# A thin-film optogenetic visual prosthesis

**DOI:** 10.1101/2023.01.31.526482

**Authors:** Eric B Knudsen, Kara Zappitelli, Jennifer Brown, Jonathan Reeder, Kevin Sean Smith, Marat Rostov, Jaebin Choi, Amy Rochford, Nate Slager, Satoru K Miura, Kyle Rodgers, Ansel Reed, Yonatan R Lewis Israeli, Seton Shiraga, Kyung Jin Seo, Corey Wolin, Paul Dawson, Mohamed Eltaeb, Arvind Dasgupta, Max Rothman, Eugene Yoon, Paul Chong, Seleipiri Charles, Jay M. Stewart, Ruwan A Silva, Tyson Kim, Yifan Kong, Alan R Mardinly, Max Hodak

**Affiliations:** Science Corporation, Alameda CA, USA; Department of Ophthalmology, University of California, San Francisco, CA, USA; Department of Ophthalmology, Stanford University School of Medicine, Palo Alto, CA, USA

## Abstract

Retinitis pigmentosa and macular degeneration lead to photoreceptor death and loss of visual perception. Despite recent progress, restorative technologies for photoreceptor degeneration remain largely unavailable. Here, we describe a novel optogenetic visual prosthesis (FlexLED) based on a combination of a thin-film retinal display and optogenetic activation of retinal ganglion cells (RGCs). The FlexLED implant is a 30 µm thin, flexible, wireless µLED display with 8,192 pixels, each with an emission area of 66 µm^2^. The display is affixed to the retinal surface, and the electronics package is mounted under the conjunctiva in the form factor of a conventional glaucoma drainage implant. In a rabbit model of photoreceptor degeneration, optical stimulation of the retina using the FlexLED elicits activity in visual cortex. This technology is readily scalable to hundreds of thousands of pixels, providing a route towards an implantable optogenetic visual prosthesis capable of generating vision by stimulating RGCs at near-cellular resolution.

## Introduction

Photoreceptor pathologies such as retinitis pigmentosa (RP) and dry macular degeneration are leading causes of irreversible vision loss, with prevalence of 0.03^1^ and 1%^2^ respectively, affecting millions of people. Although these diseases have distinct etiologies and progressions, they share a common pathology of gradual photoreceptor cell death that results in loss of vision^1, 3^. While a range of therapies to address these diseases at various stages of progression are in active development^4–10^, current clinical options are limited. In these diseases, retinal ganglion cells (RGCs) are left largely intact^11–13^. Here, we describe a prototype device with the goal of restoring vision by directly activating RGCs with a combination of an intraocularly implanted device and optogenetics.

Previous attempts at retinal prostheses can be broadly classified into two approaches: electrical stimulation and optogenetic stimulation^14–16^. Early retinal prostheses placed arrays of electrodes on the retinal surface to activate RGCs to create visual sensation^17, 18^. Electrode arrays such as the Argus II and IMIE 256 used a through-sclera via to couple retinal electrode arrays to extraocular electronics packages driven by a wearable device that provides wireless data and power^19, 20^. While these devices generate visual sensations, they suffer from poor resolution. A promising electrical stimulation approach is to implant photovoltaic electrodes in the subretinal space that simulate bipolar cells, which normally receive direct input from photoreceptors^21^. The use of photovoltaic pixels elegantly eliminates the requirement for a through-sclera via, but implanting a device between the retina and choroid requires creating a retinal detachment, which limits the area of the visual field that can be safely accessed^22^. This approach is also more susceptible to the substantial structural reorganization present in most retinal degenerative diseases^23, 24^.

Alternative approaches to visual restoration for photoreceptor pathologies aim to genetically modify RGCs to express light-gated ion channels, rendering them directly sensitive to light. Indeed, this approach to vision restoration was among the earliest proposed applications of optogenetics^25^, and has been enabled by the introduction of adeno-associated virus (AAVs) engineered to cross the inner limiting membrane (ILM) in primate retina^26^. Recently, it was shown that AAV-mediated optogenetic excitation of RGCs could drive visual cortex in primates^27–31^. A related approach to vision restoration is currently in a clinical trial, where RP patients receive an intravitreal injection of an AAV encoding the red-shifted opsin ChRimsonR^32^. Patterned optogenetic stimulation is then delivered with an extraocular light source. Using this approach, patients with minimal light perception regained some object localization^33^. However, optical stimulation at cellular resolution from outside the eye using a digital micromirror device (DMD) or spatial light modulator (SLM) must contend with significant difficulties in eye tracking and registering micromovements of the device relative to the face. Other gene therapy approaches to vision restoration seek to modify RGCs to become sensitive to ambient light levels, but these approaches must overcome the slow kinetics of these responses^34, 35^.

The diversity of coding schemes present in distinct RGC subtypes presents a problem for visual prosthetics. There are numerous classes of RGCs in the retina^36–38^, each of which carry distinct information^37, 39, 40^. For example, ON and OFF RGCs increase firing rate with increased or decreased luminance, respectively^41^. High-density multi-electrode array (MEA) recordings have spatially mapped these RGC types, and found that each uniquely contributes to the reconstruction of visual stimuli^42–44^. This indicates a mature understanding of the neural coding principles in the retina, and demonstrates that neighboring RGCs encode different aspects of visual stimuli. Therefore, an advanced visual prosthesis for photoreceptor degeneration must seek to address RGCs at cellular resolution.

Here, we propose an approach to vision restoration by optogenetic stimulation of RGCs using an implantable thin-film µLED display with pixels approximately the size of RGC somas. The flexible display is 30 µm thick and is affixed to the retinal surface using tacks. The display is connected to an electronics package mounted under the conjunctiva by a through-sclera via. We describe fabrication of the device, validation of an optogenetic viral vector, and surgical implantation in rabbits. In a rabbit model of photoreceptor degeneration, we demonstrate that optogenetic stimulation of RGCs evokes activity in the contralateral visual cortex. This approach suggests a path to an implanted optogenetic therapy that operates at cellular resolution.

## Results

### A Flexible Thin Film Optogenetic Display

We developed an implantable visual prosthesis based on a µLED display that is affixed to the retinal surface. The optical power required to activate neurons with optogenetics (>20 mW/cm^2^)^45, 46^ is easily achieved with µLEDs, which can be fabricated as small as 1 µm from gallium nitride (GaN) using conventional wafer scale processes^47^. Given the ILM is just 5-10 µm thick^48^, we reasoned that affixing the display directly to the retina could allow a single pixel to address only a few RGCs, even without collimating optics.

We fabricated a series of prototype devices in two main form factors for intraocular implantation in rabbits, the “hinge” FlexLED and the “u-turn” FlexLED (Figure 1a,b). The µLED displays are monolithically fabricated on epitaxially grown GaN-on-sapphire substrates, utilizing polyimide as the flexible backbone for multi-layer, through-via routing of primarily gold traces. Because individual µLEDs are buried in a monolithic polyimide package, they do not need to be transferred to an existing routing layer. This enables denser routing and more tightly integrated packaging layers, but comes at the cost of less efficient use of expensive epitaxially-grown GaN wafers. Each device consists of 4-6 layers of polyimide of various thickness. Electronic components are assembled directly onto the polyimide-metal structures before substrate release.

**Figure 1:**
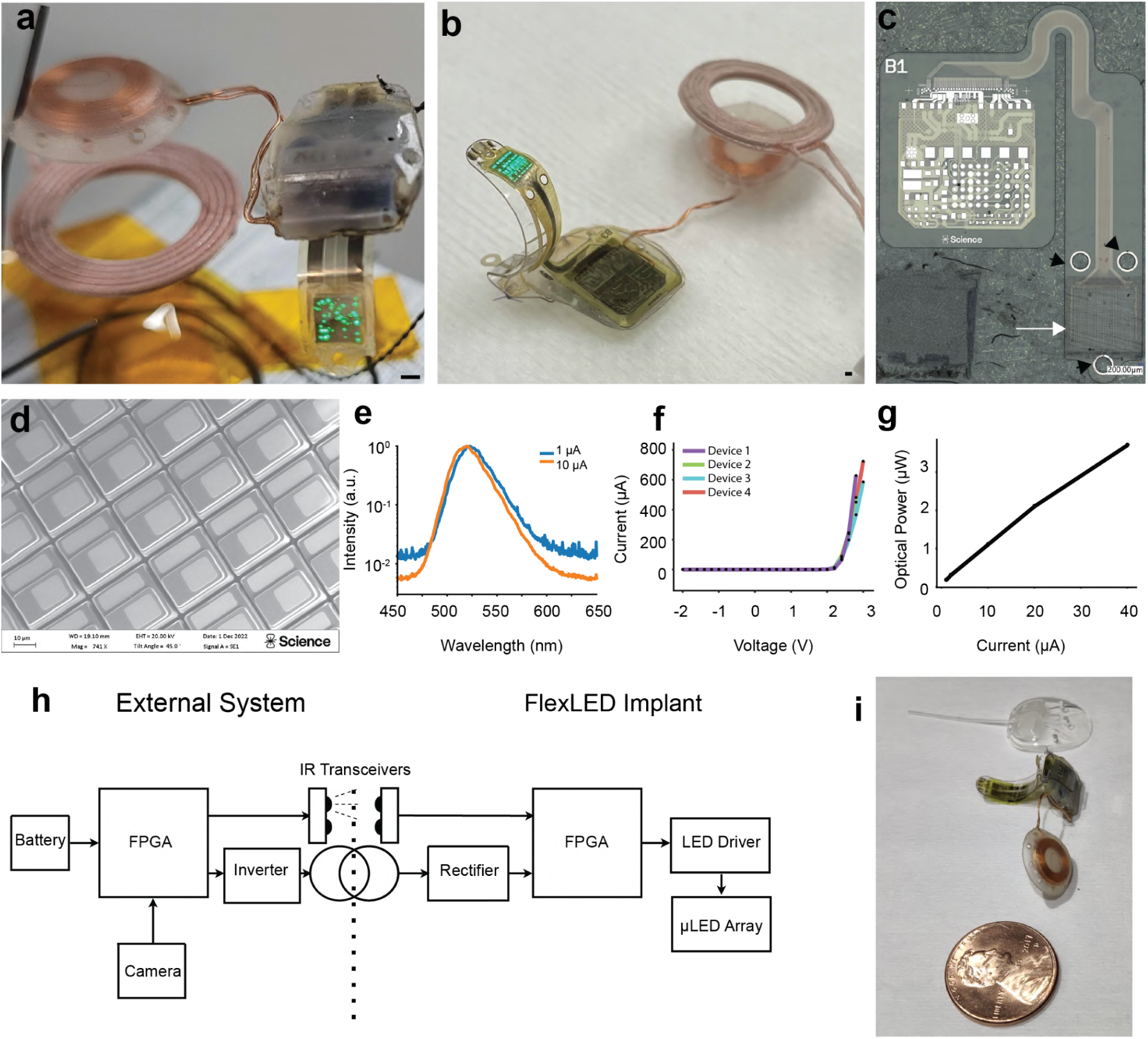
A wireless, thin-film optogenetic retinal prosthesis. a) An image of a hinge FlexLED device being wirelessly powered on a benchtop fixture and displaying a random pattern of µLEDs for illustration purposes. Black scalebar 1 mm. b) An image of a u-turn FlexLED mounted on a surgical scaffold and being wirelessly powered on a benchtop fixture and displaying a random pattern of µLEDs for illustration purposes. Black scalebar 1 mm. c) An image of the thin-film layer of a U-turn FlexLED device before bonding to the electronics package. Black arrowheads indicate holes for retinal tacks, and white arrow indicates the display area. d) SEM image of the display region of a representative hinge device. Scalebar 10 µm. e) Emission spectra of a representative hinge device at 1 µA (blue) or 10 µA (orange). f) Representative current-voltage traces for a batch of U-turn devices; the different colors indicate four unique devices under test. g) Chart showing linear relationship between drive current and optical power for a representative u-turn device. h) Block diagram showing interactions between the implanted device and external devices providing wireless power and data. i) Photograph of a hinge device with a wireless power coil next to a penny (bottom) and Ahemed valve glaucoma shunt (top). Note that the device is mounted in a rigid surgical scaffold that is removed intraoperatively.

The hinge device contains 16,000 µLED pixels with a 20 µm pitch, an active area of 66 µm^2^ (6×11 µm, Figure 1c,d), and is 30 µm thick. Due to wiring constraints, only 8,192 of the µLED pixels are connected in this prototype device. In contrast, the u-turn device contains 2,048 pixels with a 42 µm pitch, an active area of 285 or 680 µm^2^ (15×19 µm or 20×34 µm), and a thickness of 22 µm. The smaller number of µLEDs in the u-turn device decreases the number of traces that need to be routed out of the eye, eliminating mechanical reliance on an interlayer polyimide bond (the “hinge”). The hinge bond has a measured adhesive failure force of 1.2N compared to the cohesive failure strength of the u-turn device of 15.3N (Sup Figure 1).

The core considerations for µLED substrate selection were biocompatibility, availability, scalability, high efficiency, and a green emission spectrum. GaN is biocompatible^49–51^, and in the last decade has seen a surge in development as small, bright emitters are a requirement in many AR/VR applications^52–54^. GaN µLEDs scale well to small sizes due to the lower recombination velocity compared to GaP^55, 56^. While GaN µLEDs have long been known to be bright, the efficiency of GaN µLEDs has steadily improved over the last decade, allowing use in power-limited applications^57–59^. Importantly, GaN µLEDs can be pushed to green wavelengths to reduce the photochemical hazard^60^ of the device. Green wavelengths can be used to activate microbial opsins for optogenetics at light levels compatible with the photochemical hazard function while avoiding efficiency and safety issues with red µLEDs^61, 62^.

Characterization of our µLED devices are shown in Figure 1e-g. The spectral peak is at 545 nm and the full width half max is 20 nm, and exhibit the expected slight blue-shifting at higher current densities typical of green GaN^63^ (Figure 1e). Our devices were able to generate significant light output below 3V, enabling the use of low-voltage electronics (Figure 1f). We measured peak external quantum efficiency (EQE) of our uLEDs to be ∼7% by placing a 70mm^2^ photodiode (Thorlabs S120VC) against the sapphire substrate and measuring the bottom-emitted output power (Figure 1g). Since GaN-based µLED efficiency has been reported as high as 15% EQE for comparable µLEDs^47, 64^, additional process development is likely to yield further improvements in efficiency.

A block diagram of the FlexLED electronics system is shown in Figure 1h. The design is based on wireless power delivery from glasses and does not contain an implanted battery. Instead, the device is powered via a wireless coil, with onboard rectification and regulation. An IR data link allows for continuous streaming of data to and from the implant. It is capable of passing 2 Mbps and signaling the driver at up to 90 FPS, but in the absence of an active matrix, the effective operating frame rate is governed by the exposure time needed to activate the opsin. Total power consumption is an important metric in implantable devices both from the perspective of overheating tissues as well as device usability. The power consumption of the implanted device is 30-35 mW when all pixels are triggered, although this does not include efficiency losses in the wireless coil. The exact power consumption depends on the stimulation patterns, but is typically < 50 mW, including the power of the wireless coil.

We looked to glaucoma shunt devices for inspiration on form factor, as these are routinely sutured to the sclera in patients without major adverse effects or sustained discomfort^65^. The volume of the human eye is roughly 6.5 mL^66^, and a typical glaucoma shunt is < 5% of this volume, placing an upper bound on the size of a long-term ocular implant. Among the smallest of such shunt devices is the Ahmed valve^67^. The dimensions of the Ahmed shunt are roughly 16.5 x 13.1 x 2.4 mm, with a volume of ∼300 µL (as measured). In comparison, the FlexLED package which is also sutured to the eye is 11 x 10 x 2 mm with a volume of 131 µL. An additional wireless coil is attached to the FlexLED via a flexible tether allowing for free positioning, with a 10 mm diameter, 1.5 mm height, and volume of 119 µL (Figure 1i). The FlexLED electronics are encapsulated with an elastomeric polyurethane cap in addition to an epoxy underfill to provide more patient comfort. A 2-3 µm layer of Parylene-C encapsulates the full device to provide a high lubricity and abrasion resistant layer during handling.

**Table 1:** Device Specifications

| Specification | U-Turn Device | Hinge Device |
| --- | --- | --- |
| Display Area | 2.7 mm x 2.7 mm | 2.56 mm x 2.56 mm |
| Total Number of Pixels | 4,096 | 16,000 |
| Number of connected pixels | 2,048 | 8,192 |
| Per-pixel emission area | 285 - 680 μm<sup>2</sup> | 66 μm<sup>2</sup> |
| Center wavelength | 545 nm | 545 nm |
| Maximum Framerate | 90 FPS | 90 FPS |
| Maximum Bandwidth | 2 Mbps | 2 Mbps |
| Receive Latency | < 1 ms | < 1 ms |
| Power Consumption* | 30 mW | 35 mW |
| Volume of package | 144 μL | 131 μL |
| Volume of wireless coil | 119 μL | 119 μL |
| Thickness of display area | 22 μm | 30 μm |

\* not including efficiency losses from wireless coil

### *In vitro* validation of optogenetic viral vector

Taking into account the optical power available from the FlexLED and the photochemical power function, we generated a viral construct that would allow robust optogenetic activation of RGCs with brief pulses. Since the passive display drivers on the prototype FlexLED allow illumination of one row at a time, brief illumination times are required to achieve high frame rates (64 rows require ∼0.5 ms for 30 FPS). For the opsin, we chose ChRmine-mScarlet^46^, a fluorescently-tagged light-gated optogenetic activator, whose expression was placed under the control of a promoter derived from the human Synapsin1 (hSyn1) gene. The opsin additionally contained a soma-targeting sequence derived from the potassium channel Kv2.1^68^. The construct was packaged in AAV2.7m8, a variant of synthetic AAV vectors engineered to cross the ILM^26^.

To validate the opsin, we first transduced DIV14 iPSC-derived human neurons with the construct and performed whole-cell electrophysiological recordings during optical stimulation. ChRmine-expressing human neurons were anatomically normal, hyperpolarized at rest, and fired action potentials in response to current injection (Figure 2a-b). All neurons with fluorescence exhibited light-evoked photocurrents with kinetics characteristic of ChRmine^46^, and most fired light-evoked action potentials (Figure 2b). We recorded in both voltage and current clamp configurations while delivering light stimuli of varying duration and intensity (Figure 2c, center wavelength 565 nm). Stimuli as short as 0.5 ms could elicit the occasional action potentials, but 5 ms stimuli above 7 mW/cm^2^ were required to elicit action potential with ≥90% probability (Figure 2c, mean of n=10 neurons). These stimuli elicited photocurrents with amplitude −0.952±0.57 nA (mean±s.d.) which was less than half of the maximal photocurrents recorded (−1.68±0.57 nA mean±s.d., Figure 2c, data represent mean values from n=10 neurons). These data indicated that pulse durations between 1 and 5 ms are needed to reliably activate these cells with irradiances that are safe for long-term exposure in the retina. This bounds the frame rates we can achieve using a passive display.

**Figure 2:**
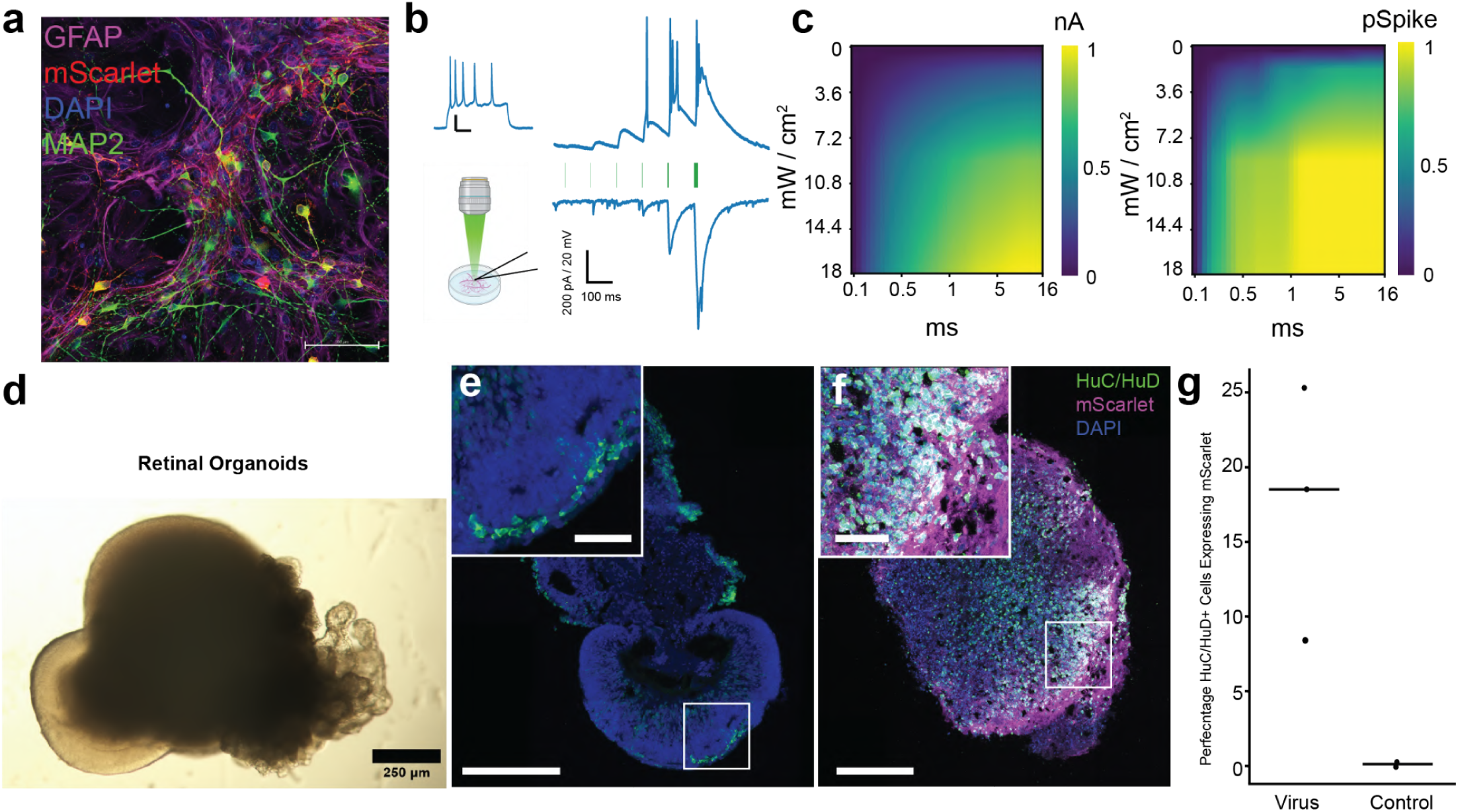
*In vitro* characterization of ChRmine-mScarlet-Kv2.1 in human cells. a) Confocal microscope image of cultured human neurons derived from IPSC on a mouse glia monolayer (DIV35, GFAP: magenta, mScarlet: red, MAP2: green, DAPI: blue, scalebar 100 µm). b) Whole cell patch clamp recording of virus-transduced human neurons derived from iPSCs. Neurons fired action potentials upon current injection (representative trace, top, scalebar 100 ms, 20 mV). Representative traces from a neuron undergoing optical stimulation in current clamp (top) and voltage clamp (bottom) are on the right (pulse durations of 0.1, 0.2, 0.5, 1, 5, 16 ms, 18 mW/cm^2^). Scalebar 100 ms, 200 pA (voltage clamp) or 20 mV (current clamp). c) Mean data from n=10 neurons in voltage clamp (left) or current clamp (right) showing the peak light-evoked photocurrents amplitudes or spike probability (colorbar) as a function of exposure duration (x-axis) or irradiance (y-axis). d) Representative light microscope image of a human retinal organoid (Day 125, scalebar 250 µm). e) Confocal image from a cryosection of a D65 control organoid stained with HuC/HuD (green) and DAPI (blue). Magenta (not visible) shows background mScarlet signal. Scalebar 250 µm; inset 50 µm. f) Confocal image from a cryosection of a D65 organoid exposed to 5e10 vg of hSyn1-ChRmine-mScarlet-Kv2.1 (magenta) fixed 28 days later, and stained for HuC/HuD (green) and DAPI (blue). Scalebar 250 µm; inset 50 µm. g) Quantification of HuC/HuD+ cells expressing mScarlet

Next, we verified that the viral construct could express the opsin in human RGCs. To this end, we generated human retinal organoids from iPSCs, transduced them with virus, and analyzed the expression of ChRmine-mScarlet via histology (Figure 2d-g). 65-day-old organoids robustly expressed HuC/HuD, a commonly used marker for RGCs in human retinal organoids^69–72^. Unlike in the retina, where RGCs are located close to the surface, these HuC/HuD+ cells were located throughout the organoid. Quantification of mScarlet-expressing cells reveals that ∼20% of HuC/HuD+ cells were transduced, and cells on the surface of the organoid showed increased transduction rates (Fig 2e-f). The ectopic location of putative RGCs deep inside the organoid accounts for the relatively low rate of transduction. This suggests that the viral vector can transduce human RGCs and express ChRmine-mScarlet.

### *In vivo* validation of optogenetic viral vector

We decided to perform *in vivo* validation of our FlexLED-optogenetic approach in New Zealand white rabbits. This required that, prior to implanting the FlexLED in the eye, we 1) establish a method to record retina-driven responses in visual cortex, 2) establish a rabbit model of photoreceptor degeneration, 3) confirm that our AAV construct is capable of transducing RGCs in the retina, and 4) confirm that activation of the opsin in the RGCs can evoke responses in visual cortex.

To establish a method to record and characterize retinal-evoked responses, we chronically implanted electrocorticography (ECoG) grids in the primary visual cortex and began by establishing retinotopic maps through visually-evoked activity. The grids covered an area of 4 mm mediolateral by 8 mm anteroposterior and sampled surface potentials at a 1 mm pitch from 200 µm electrodes (Figure 3a). Under general anesthesia, we presented high contrast visual stimuli of 10 degrees in visual angle to the contralateral eye (spherically corrected based on distance of the eye from the screen, typically 10-20 cm) over a visual field of 80 degrees horizontal by 60 vertical. The stimuli reliably elicited visual-evoked activity, confirming the ECoG grids covered the visual cortex (Figure 3b). Responses were highly confined in cortical space: a median of 3 electrodes (spanning about 2-3 mm, interquartile range 1-9) exhibited significant responses to these stimuli, with a strong tendency for neighboring electrodes to be coactive (Figure 3c-d). Responses could be decoded across a large portion of the visual field, suggesting we have access to a retinotopic map (Figure 3e-f). This map suggests that optogenetic activation of a retinal region corresponding to 10 degrees of visual angle should be as spatially confined as these visual stimuli.

**Figure 3:**
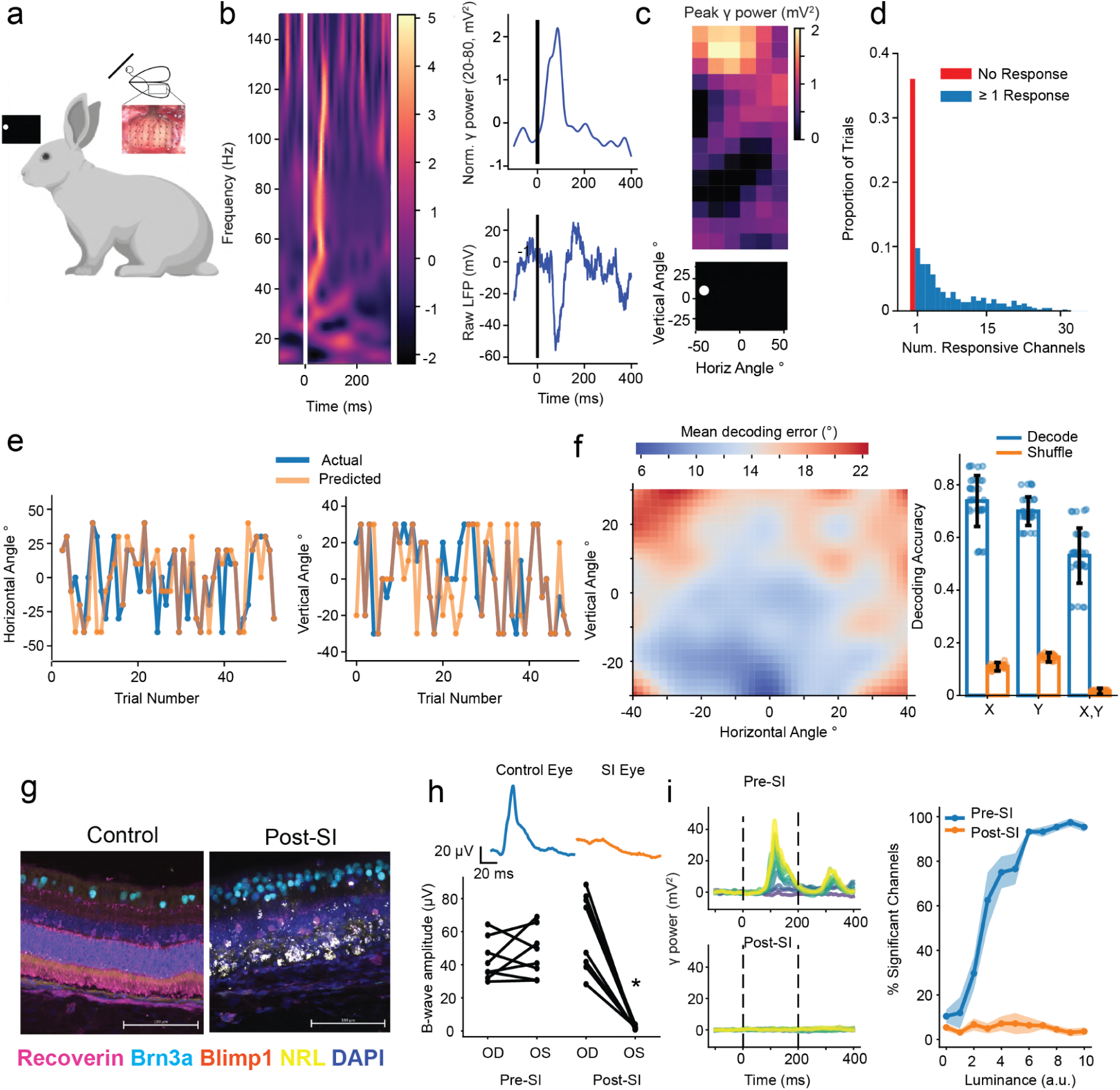
Chronic recording from Rabbit visual cortex in a model of photoreceptor degeneration. a) Schematic of experimental approach: New Zealand white rabbits are implanted with a chronic 32-channel ECoG grid over visual cortex, and visually-evoked potentials are recorded as high contrast stimuli are presented using a monitor. b) Left, example single trial spectrogram showing the power (colorbar, mV^2^) as a function of frequency (y-axis) over time after presentation of a visual stimulus. Bottom right, raw LFP (mV) versus time from stimulus presentation. Top right, normalized gamma power over time from stimulus presentation. c) Representative single trial data showing focal activation of the ECoG grid by a high contrast visual stimulus of 10 degrees visual angle. Top, heatmap showing peak gamma power during stimulus presentation. Bottom, depiction of the visual stimulus. d) Histogram showing the number of significantly responding ECoG channels (max 32) for each trial. Trials where no channels were significantly modulated are shown in red (median 3 modulated channels, interquartile range 1-9 channels). Data represent 11025 trials from 7 rabbits (1575 trials per rabbit). e) Actual (blue) versus predicted (orange) horizontal (left) or vertical (right) visual angle over the duration of a representative 50-trial recording session. f) Left, heatmap plotting the mean decoding error in degrees as a function of the horizontal and vertical visual angle of the presented stimulus (see colorbar). Right, plot showing decoding accuracy for horizontal (x), vertical (y), or both (x,y) across all recording sessions (blue) and with stimuli labels shuffled (orange). Data represent 7 recording sessions from 7 rabbits. g) Confocal micrographs from rabbit retinas injected with sodium iodate for 6 weeks (right) or controls (left). Images were obtained from 25 µm cryosections and are maximum intensity projections from z-stacks with 0.5 µm slices. Images are Recoverin (Magenta), Brn3a (Cyan), Blimp1 (Orange), NRL (Yellow) and DAPI (blue). Scale bars are 100 µm h) Top, representative traces from an electroretinogram recording from a control eye (blue) or sodium-iodate injected eye (orange). Scale bars: 20 µV, 20 ms. Bottom, chart showing B-wave amplitude for control eye (OD) or SI-injected eye (OS) either before (left) or > 4 weeks after sodium iodate injection (right). * indicates p < 0.01, paired t-test. i) Left, gamma power (20-80 Hz, mV^2^) from a representative animal showing full-field stimuli of increasing luminance (colors) before (top) or after (bottom) injection of sodium iodate into the contralateral eye. Right, plot showing % of significantly responding ECoG channels before (blue) or after (orange) to full-field visual stimuli of increasing luminance. Data are mean and s.e.m. from 6 rabbits pre-SI, 7 post.

Next, we generated a rabbit model of photoreceptor degeneration by intravitreal injection of sodium iodate^73^. We first confirmed this paradigm in mice^74^ (Sup Figure 2a). Histological examination of rabbit retinas 6 weeks after injection confirmed widespread loss of photoreceptor outer segments and general disruption of the inner retina, but with little impact on RGCs (Figure 3g). Degeneration of the photoreceptors were also functionally confirmed by a significant reduction in the amplitude of the photopic electroretinogram (ERG) B-wave two weeks after sodium iodate injection (Figure 3h). Luminance responses in the contralateral visual cortex were also strongly attenuated, further confirming loss of photoreceptor function (Figure 3i). Thus, this chemical model mimics the photoreceptor degeneration diseases that are likely to benefit from a visual prosthesis.

Having established a paradigm to record from the visual cortex in the context of a degenerating retina, we evaluated our AAV’s expression in the rabbit retina. We first confirmed that the AAV could cross the ILM to transduce RGCs via intravitreal injection in mice (Sup Figure 2c-d). We then evaluated the expression of ChRmine-mScarlet in the rabbit retina after intravitreal injection (1e11 vg/eye). Histological examination of retinas 6 weeks after injection of virus showed widespread and robust transduction of RGCs, based on their anatomical position in the retina and co-labeling with RNA-binding protein with multiple splicing (RBPMS), an RGC-specific marker^75^ (Figure 4a). We further analyzed virus expression in rabbit retina via scRNA-seq, as we did for our retinal organoids. scRNA-seq protocols recovered all major cell types from the retina, and we found strong selectivity of ChRmine-mScarlet in RGCs compared to all other classes of retinal cells (Sup Figure 3a-b). Although sequencing indicated that only ∼20% of RGCs expressed ChRmine-mScarlet, a similar percentage expressed synapsin1 mRNAs. Since read counts were generally low in these experiments (average 1.64±0.66 for ChRmine-mScarlet, 0.97±0.32 for synapsin1, normalized counts +/- s.d.; *n* = 69 and 56 cells, respectively), it is likely this method significantly underestimates the true proportion of positive cells due to a large number of drop-outs. Indeed, imputation analysis suggests ∼90% of RGCs could be expressing the virus (377 out of 417 RGCs; Methods). Importantly, ERG recordings from eyes injected with virus, but not sodium iodate, revealed no changes in visual responses (Sup Figure 4a-e). Furthermore, immunostaining of virus-injected retinas did not indicate any inflammation or immune cell infiltration (Sup Figure 4e-f). Taken together, the above data indicate that our AAV construct efficiently and specifically transduces RGCs.

**Figure 4:**
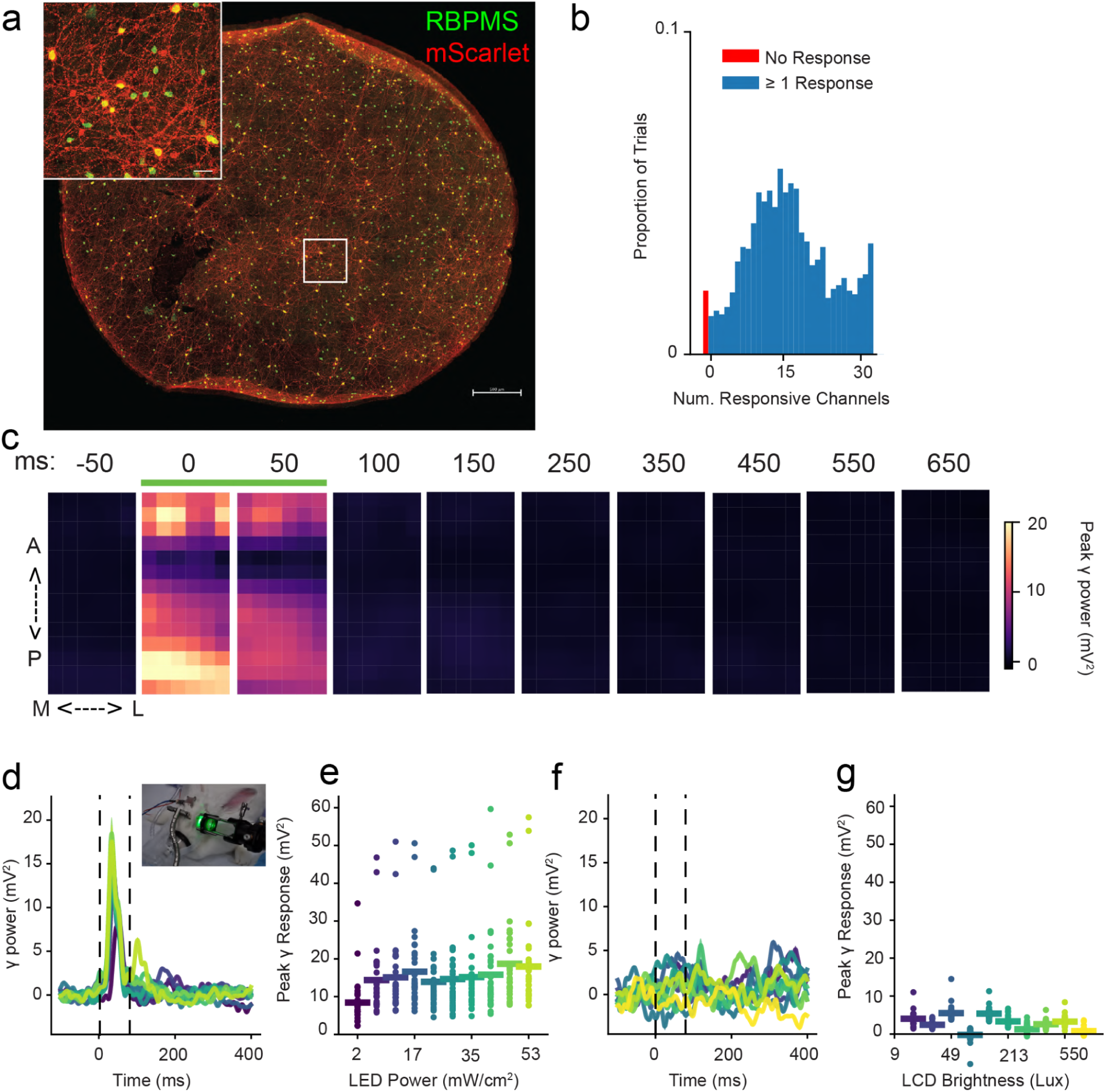
Optogenetic activation of V1 by hSyn1-ChRmine-mScarlet-Kv2.1 in RGCs. a) Representative flat mount of rabbit retina expressing hSyn1-ChRmine-mScarlet-Kv2.1 (red) and stained for RBPMS (green). Scale bar: 500 µm, inset: 50 µm. b) Histogram showing the number of significantly responding ECoG channels (max 32) for each trial of optogenetic stimulation using a 530 nm LED. Trials where no channels were significantly modulated are shown in red (median 16 modulated channels, interquartile range 11-22 channels). c) Series of heatmaps showing peak gamma power (20-80 Hz, mV^2^, see colorbar) as a function of time from stimulus onset (top, green bar), versus position on the grid (M-L: medial, lateral; A-P: anterior, posterior). Data are the mean of 550 trials from a representative of n = 4 Rabbits. d) Mean responses of all ECoG channels showing peak gamma power (20-80 Hz, mV^2^) versus time for optogenetic stimuli of increasing intensity (colors). Dashed bars indicate stimulus timing. Inset, image of experimental setup. e) Peak gamma response for single trials (dots) and trial means (bars) as a function of LED power measured outside the eye (mW / cm^2^, x-axis). Data show n = 550 trials from a representative of n = 4 rabbits. f) Mean responses of all ECoG channels showing peak gamma power (20-80 Hz, mV^2^) versus time for visual stimuli of increasing intensity (colors). Dashed bars indicate stimulus timing. g) Peak gamma response for single trials (dots) and trial means (bars) as a function of LCD monitor brightness (Lux, x-axis). Data show n = 330 trials from a representative of n = 4 rabbits.

We next tested the ability of extraocular LED stimulation to drive optogenetically-evoked activity in visual cortex for eyes that were injected with both virus and sodium iodate. As expected, high contrast full-field visual stimuli from a computer monitor failed to evoke significant responses (Figure 4f-g). In contrast, full-field excitation using a 530 nm LED light elicited responses across the ECoG grid (Figure 4b-e), with a median of 16 channels (11-22 interquartile range) displaying significant responses (Figure 4b). This wide-spread activation of visual cortex is distinct from ECoG activation patterns generated by visual-evoked experiments (Figure 3d), and likely reflects activation of opsin-expressing RGCs distributed widely in the retina. Thus, the data suggest that we are able to optically evoke activity in the retinal cells transduced by our AAV construct.

### Modeling FlexLED optical stimulation

Having validated the rabbit optogenetic preparation, we considered how the FlexLED could stimulate the retina. We first assessed whether our vector was compatible with radiative exposure safety considerations for a long-term implanted device. Damage to the retina can occur via either photochemical or thermal processes; we calculated that for a device in contact with the retina, the long-term potential for damage arises primarily from photochemical effects, rather than thermal considerations (Sup Fig 5). Taking into consideration the spectral dependence of the aphakic photochemical hazard function^60^, the maximum peak irradiance allowed for chronic epiretinal µLED centered at 545 nm and operating continuously is 70 mW/cm^2^ (Sup Fig 5). This is well below the threshold for neural activation measured *in vitro* (Figure 2a-c), even for a worst case scenario of constant illumination.

We next considered the spatial resolution we could achieve using the FlexLED. Since the device does not contain electrodes and is not optically transparent, recording directly from RGCs *in vivo* was not possible. Instead, we modeled the response of RGCs to FlexLED stimulation. We first experimentally determined the angular emission profile from single FlexLED pixels in a fluorescent dye solution (AF568, 10 µM) and using a goniophotometer and found that it closely resembled a Lambertian distribution (Sup Figure 6a-b). We then used these empirically determined profiles to perform Monte Carlo modeling of light scattering and absorption inside the retina created by each of the hinge and u-turn devices (Sup Figure 6c-d). We calculated that for a µLED optical power of 2 µW, a value achievable with < 20 µA (Figure 1g), a RGC aligned with a µLED center would see an irradiance of ∼30 mW/cm^2^, an optical power that should elicit reliable action potentials and is well below the safety limit for continual irradiance.

We then used the physical layout of µLEDs and the location of RGCs measured from peripheral rabbit retina to calculate the mean irradiance on each RGC soma from each pixel (Sup Figure 6e). We passed this irradiance through an action potential threshold obtained from *in vitro* recordings and repeated the calculations for a variety of optical powers, RGC activation field sizes (soma and proximal dendrites), and axial distances between the µLED pixels and RGCs. To account for the thickness of the ILM and axon tracks, we assumed RGCs were located at least 20 µm from the surface of the retina. These simulations showed that if the µLED array could be secured < 50 µm away from the RGCs, and assuming that all RGCs express opsin, almost every RGC located under the display area could be activated by at least one µLED pixel (Sup Fig 6f). We next calculated the number of pixels that could activate any addressable RGC. Across a range of conditions, on average most RGCs could be activated by just 1 or 2 µLEDs (mean ∼1.5). This indicates that the device should be capable of operating at cellular resolution in the peripheral retina (Sup Fig 6g). At high powers, up to ∼50% of pixels could activate at least one RGC (Sup Fig 6h). Notably, the size of the RGC activation field did not significantly impact results, and there were no large differences in the simulation results between the hinge and u-turn devices. This indicates that even the u-turn device, with its larger and less dense µLED pixels, may oversample RGCs in peripheral rabbit retina. Notably, simultaneous activation of the full array should activate nearly all the RGCs located under the display at lower powers and at considerably farther axial distance (Sup Fig 6i).

### FlexLED-evoked responses in visual cortex

We implanted the FlexLED device in two animals where we observed cortical responses to extra-ocular optogenetic stimulation in the context of degenerated photoreceptors. To support surgical implantation, the FlexLED device was mounted on a rigid surgical scaffold that takes advantage of the tack-hole near the display. The scaffold is curved so that the active area is positioned towards the anterior chamber after insertion, mitigating the possibility of contacting the retina while the package is sutured to the globe. The scaffold features a “rip-cord” (Figure 5a) that allows the surgeon to release it while the active area is inside the eye by pulling on a tab outside the eye. The rest of the surgical procedure is based on standard retinal surgery techniques (see methods for detailed procedure). Briefly, the eye is dilated, and a partial 180° peritomy is performed. A phacoemulsification is performed to remove the lens, followed by a core vitrectomy. The FlexLED, supported by the surgical scaffold, is inserted through a 5 mm scleral incision, and the extraocular electronic package is secured to the sclera via non-absorbable sutures (Figure 5b, top). The FlexLED carrier is removed, the sclerotomy sealed, and intraocular forceps are used to grip a tab placed on the back of the FlexLED array. A tack is used to secure the display to the retina (Figure 6b, bottom). The eye is sealed, the FlexLED power coil is tucked next to the device, and the conjunctiva is sutured. Figure 6c shows a control eye (top) and an implanted eye (bottom) 4 days after surgery.

**Figure 5:**
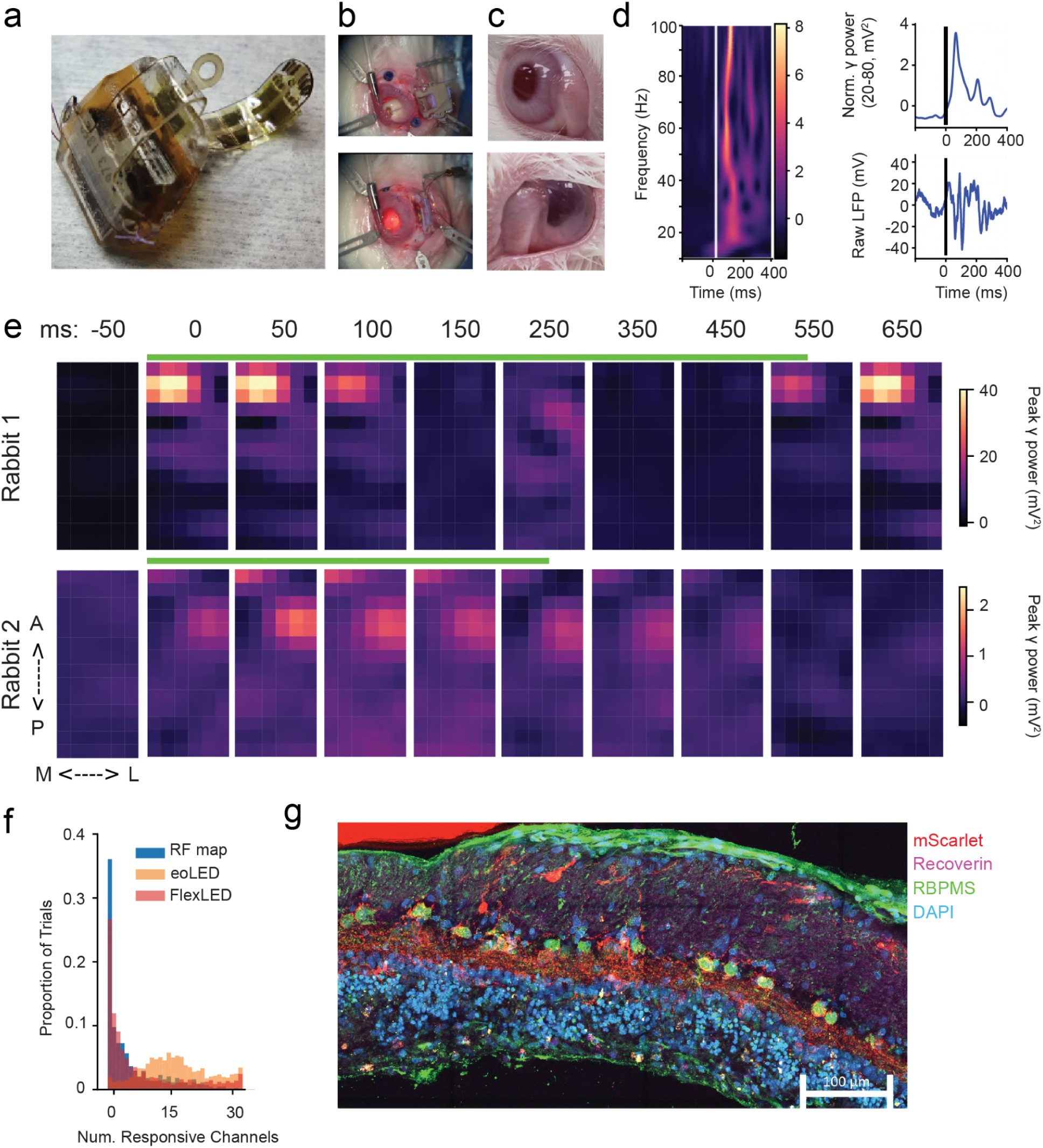
A thin-film optogenetic visual prosthesis drives visual cortex in a pattern that resembles visual responses. a) An image of a hinge version of the FlexLED in a surgical scaffold b) Top, surgical image showing the FlexLED sitting on the globe as the array, still attached to the carrier, is inserted through a sclerotomy. The carrier holding the array is visible in the anterior chamber, protecting it from hitting the retina. Bottom, the FlexLED package is secured to the watertight eye, and a tack is prepared to secure the device to the retina. c) Close up images of the subject’s eyes just four days after the procedure. Top, control eye, bottom, implanted FlexLED eye (note that the lens has been removed). d) Left, example spectrogram showing the power (colorbar, mV^2^) as a function of frequency (y-axis) over time after presentation of an optogenetic stimulus through the FlexLED. Bottom right, raw LFP (mV) versus time from stimulus presentation. Top right, normalized gamma power over time from stimulus presentation. e) Series of heatmaps showing peak gamma power (20-80 Hz, mV^2^, see colorbar) as a function of time from FlexLED stimulus onset (top, green bar), versus position on the grid (M-L: medial, lateral; A-P: anterior, posterior). Data are the mean of 800 trials from N = 2 rabbits (each row corresponds to one rabbit). f) Histogram showing the number of significantly responding ECoG channels (max 32) for each trial of FlexLED optogenetic stimulation (red, median 4 modulated channels, interquartile range 1-12 channels). Data are shown superimposed on the histograms from Figure 3d (beige, extraocular optogenetic stimulus, and Figure 4b (blue, high-contrast 10 degree visual stimulus). g) Confocal microscope image from a section of retina obtained from directly beneath the active area of the device for Rabbit 1 in Fig 6e stained for mScarlet (Red), RBPMS (green), Recoverin (Magenta), and DAPI (blue). Note that a previous sodium iodate injection has ablated photoreceptors and has resulted in disorganization of the inner retina. Scalebar 100 µm.

After implantation, we used the FlexLED to drive AAV-transduced cells and evoke activity in the contralateral visual cortex. Although several lines of evidence indicated that sodium iodate ablated photoreceptors and visual responses, to rule out contributions from any remaining photoreceptors, we performed these experiments after intravitreal injection of a cocktail of synaptic blockers. We verified the efficacy of the blockers by observing that they completely blocked light-evoked responses in naive animals (Sup Figure 7a). The prototype devices we used to collect this data had suboptimal pixel yield and exhibited pixel-to-pixel variance in brightness, so we focused on full-array activation of the FlexLED device at high contrast. The FlexLED evoked an obvious and reliable visual cortex response (Figure 5d). Like the natural visual response to localized high contrast stimuli, but in striking contrast to full-field extra-ocular optogenetic stimulation, stimulation via the FlexLED resulted in focal visual cortex responses in both subjects (Figure 5e-f). These responses were spatially constrained to a small number of electrodes (median: 4, interquartile range: 1-12), exhibited a latency of ∼50 ms, and were active following cessation of the stimulation (Figure 5e-f). In one subject, we obtained histology from the area of tissue directly under the FlexLED, and confirmed robust virus expression and degeneration of photoreceptors in that region. These data offer proof of concept that an implanted optogenetic retinal display can work in tandem with a viral construct to drive robust activity in visual cortex in a rabbit model of photoreceptor degeneration.

## Discussion

In this work, we describe the design and synthesis of a prototype retinal implant, validate a viral vector, and demonstrate that the device can drive visual cortex activity in a rabbit model of photoreceptor degeneration. The device has an active area of 2.56 x 2.56 mm, corresponding to ∼10 degrees of visual space in humans^76^. Importantly, the patterns of FlexLED-evoked activity resembled activity elicited by high-contrast 10 degree stimuli much more than those from full-field optogenetic stimulation from outside the eye (Figure 6f). This suggests that we have achieved targeted stimulation of the retina using the FlexLED device.

The FlexLED device described in this study has 8,192 pixels at up to 90 FPS at 8-bit global intensity levels. In comparison, no previous retinal stimulation device of which we are aware has achieved this pixel density^20, 22^ (1250 px/mm^2^). This version of the FlexLED uses a passive driver, which limits frame rates when taken in conjunction with the minimum times required to reliably elicit optogenetically-evoked action potentials at safe light levels. This can be remedied in the future by using active pixels in the display, which will require the addition of a thin-film transistor backplane on the device. While the number of pixels in these prototype devices is nearly two orders of magnitude below the number of RGCs in the human eye (1.2 million)^77^, wafer scale fabrication processes can be readily applied to create megapixel devices. Modeling suggests that increasing the density of µLEDs beyond the characteristic spacing of RGCs somas is unlikely to affect resolution in peripheral retina, but increasing the area on the retina will grant access to more of the visual field. Surgical constraints on the scleral incision size will likely require larger displays to be unfolded inside the eye during surgery. An active display with more pixels will also require driver electronics to be placed inside the eye. This would allow for a large reduction in the number of wires routed through the sclera, which could in turn allow more flexible positioning of the device.

The FlexLED device in this study is intended to achieve optogenetic control of the peripheral retina using a display with stimulation pixels roughly matched in size and pitch to the density of RGCs. Achieving single-cell resolution with 1-photon optogenetic excitation *in vivo* is usually extremely challenging without complex optics because of significant axial light propagation^78^. This problem of axial resolution is obviated in the peripheral retina, where RGCs are arrayed in a monolayer^79^. In the perifoveal region, where RGCs are stacked in layers up to 10 cells deep, axial light propagation may decrease resolution, but sparse viral transfection could limit the number of off-target RGC activations. Although the operating principle of this device, supported by modeling, suggests that each pixel should activate at most 2 RGCs, we did not directly measure the spatial resolution of optogenetic stimulation with the FlexLED in this study. Due to the device being secured to the retina’s surface, simultaneous stimulation and measurement of RGC activity, either via MEA recording or genetically encoded calcium indicators, poses significant technical challenges.

The resolution of the device is also influenced by the density of viral transduction. A performant viral vector for an optogenetic prosthesis must have a high transduction rate, express sufficient protein to allow activation of RGCs, and exhibit low immunogenicity. Most protocols to virally transduce RGCs require a single intravitreal injection^33^, but the virus may only persist in the eye for several days^80^, limiting its ability to achieve widespread coverage. Viral vectors have been increasingly deployed to the eye^81^ due to the notion that the eye is an “immune privileged”^82^ compartment, yet it may still lead to an inflammatory response^83^. In our work, we observed no clinically relevant inflammation resulting from our viral vectors, and histology did not reveal any infiltration of immune cells from the choroid to the retina. However, our endpoints were relatively short, and the long term potential for opsin immunogenicity remains unknown.

More generally, the risk of an immune response to the microbial opsin ought to be proportional to the number of cells that present the microbial opsin in HLA complexes, as each additional opsin-expressing cell increases the likelihood of an immunogenic interaction with the immune system. Paradoxically, optogenetic therapies should thus seek to minimize the number of opsin-expressing cells. One way to accomplish this is to encode cell-type specific expression by choice of the promoter. Although the PIONEER study employs a CAG promoter^33^, which should express opsin in most transduced cells, we elected to use a human synapsin promoter^84^, which should confine opsin expression to neuronal cells. In our study, this approach resulted in robust and selective expression of ChRmine-mScarlet in RGCs in mice, rabbits, and human organoids. At least one report suggests that this promoter may not be effective in Rhesus macaques^85^, so further study in primates is required to assess the suitability of this vector for clinical translation, although this promoter has been used successfully in primate brain^86–88^. Another way to limit the number of opsin-expressing cells is to spatially restrict the virus in the retina. Since the implant covers a narrow field of view on the retina, in the case of an intravitreal virus injection, most of the opsin-expressing cells are ‘wasted’ as they are not optically accessible to our device. In the future, embedding the virus on the display surface, for instance with a silk fibroin hydrogel^89^, could dramatically decrease the number of opsin-expressing cells in the eye without changing the function of the device.

An implanted retinal display has several advantages compared to optical stimulation from outside the eye. Stimulation from goggles must contend with both macro and microscopic movement of the goggles relative to the eye, rapid changes in eye position, and variable pupil diameter, which affects the numerical aperture of the system. Stimulating RGCs with single-cell precision from outside the eye requires extremely low-latency closed-loop tracking and computation, which is difficult to implement. In contrast, an implanted display features a fixed mapping between a given pixel and a specific RGC or set of RGCs. The display moves with the eye, eliminating the need to dynamically correct stimulation patterns. While eye tracking is still required to render a scene on the implant, the stimulation pattern does not need to dynamically track a moving target. Indeed, adding an inertial measurement unit on the implant could greatly facilitate tracking an implanted eye.

An implanted display is also fundamentally more efficient in terms of pixel use: for an augmented reality display, a visual scene must be rendered in high definition across the entire field of view, since the eye may saccade to any part of the visual scene. In contrast, an implanted display’s resolution can degrade concentrically away from the fovea to match the density of RGCs. This distinction is critical for designing an implant that takes advantage of retinal coding principles: a display that renders an entire visual scene cannot know before a saccade which RGCs it will address, so the encoding potential of the prosthesis is limited. Finally, a stable mapping of pixels to cells using an implanted display may help with plasticity^90^ even in the case of poor encoding, since downstream structures will receive consistently patterned input.

Leveraging knowledge of retinal coding schemes requires methods of calibrating stimulation. This has been achieved *ex vivo* using high density MEAs to map RGC subtypes based on electronic properties and receptive fields, followed by careful microstimulation^44, 91–93^. Future FlexLED devices could integrate recording electrodes with optogenetic stimulation to allow an *in vivo* mapping and calibration procedure to help identify RGC types and thereby calibrate the device. Patient reports of the perceptual consequences of stimulation may also serve as the basis for calibrating an encoding scheme. While the perceptual consequences of stimulating single RGCs in humans are not known, humans are able to detect single cone stimulation^94^. Since cones often impinge on a single midget ganglion cell^95^, patients may be able to consciously perceive single RGC stimulation and report on the location and affect of the sensation.

The data in this study do not directly address the question of whether the FlexLED can restore any degree of visual perception. While evoked potentials in visual cortex are highly suggestive of visual perception^96^, recordings were performed under general anesthesia. Further work in awake, behaving animals is needed to address whether and how the FlexLED drives visual perception, and experiments using recording techniques with single-cell resolution will address the functional resolution and bitrate of these optogenetic implants. Experiments in foveated animals performing psychophysical behavior during electrophysiological recordings will be informative for both of these questions. Future studies must also address device longevity and performance for chronic implantation. Although numerous challenges remain, this work demonstrates a proof-of-concept approach to optogenetic vision restoration that could in principle scale to control most of the RGCs in a human eye at near-cellular resolution.

## Methods

### Cell Culture

Primary mouse glia was obtained from C57/B6J mice on postnatal day 1-2 as described^97^. Briefly, brains were isolated and the cortices were removed after carefully stripping away the meninges with forceps. Cortices were dissociated in 1mL of 0.25% Trypsin-EDTA (1x, Gibco) and 1% DNase (Worthington) at 37°C for 6 minutes at which point DMEM (Gibco) was added. Cells were then centrifuged at 300 RCF for 5 minutes, resuspended in DMEM media with 10µM Y-27632 (ROCK inhibitor, Tocris), and plated. Primary glia were cultured in DMEM supplemented with 10% FBS (Sigma-Aldrich), 1% Penicillin-Streptomycin (100X, Gibco), and 2mM Glutamax (100X, Gibco). Stocks were maintained on 6-well tissue culture treated plates (CELLTREAT) coated with 1% Geltrex (Gibco) or glass coverslips coated with 0.01% w/v poly-d-lysine (Gibco) and 0.01% w/v laminin (Sigma-Aldrich).

Differentiation of induced pluripotent stem cells to cortical neurons was performed as described^98^ with minor modifications. Briefly, IP11NA stem cells (Allstem) were plated at a density of 30,000 cells/cm^2^ in mTesR+ media (STEMCELL) containing 10µM Y-27632 on Geltrex treated 6-well tissue culture treated plates. Cells were maintained until they reached 60-70% confluence. For days 1-3, cells were differentiated in N2 media, containing DMEM/F-12 (Gibco) with 2mM Glutamax (100X, Gibco), 16.4mM D-glucose solution (Sigma), and 1% N2 supplement (Gibco). On day 1, the media was supplemented with 10µM SB-431542 (R&D Systems), 2µM XAV-939 (STEMCELL), 100nM LDN-193189 (STEMCELL), and 2µM doxycycline hyclate (R&D Systems) which was maintained throughout the differentiation. Day 2 media contained 5µM SB-431542, 1µM XAV-939 and 50nM LDN-193189. Day 3 media was not supplemented apart from the doxycycline. From day 4 onward, the cells were cultured and matured in Neurobasal media (Gibco) with 2mM 100X Glutamax, 16.4mM Glucose, and 0.5% MEM Non-Essential Amino Acids (10X, Corning), which was supplemented with 2% B27 plus (50X, Gibco), 20nM Brain-Derived Neurotrophic Factor (BDNF, PeproTech), 20nM Ciliary Neurotrophic Factor (CNTF,PeproTech), and 20nM Glial Cell-Derived Neurotrophic Factor (GDNF,PeproTech). From day 5 onward, 5µM Ara-C (Sigma) was added. Cells were dissociated in ACCUTASE (STEMCELL) between days 5-7 and plated onto primary mouse glia monolayers at a density of 10,000-20,000 neurons/cm^2^.

5e^9^ viral genomes of AAV2.7m8 hSyn1-ChRmine-Kv2.1-WPRE were applied to each coverslip of differentiated neurons at DIV12 after a 50% media exchange, for an estimated MOI of 5e^4^. The virus was not washed off, but slowly removed by successive 50% media changes every other day.

### Whole Cell Electrophysiology

Human neuron co-cultures grown on glass coverslips were transferred from the incubator to a submerged recording chamber maintained at 20–25 °C by heating in a solution containing (in mM) NaCl 119, KCl 2.5, MgSO4 1.3, NaH2PO4 1.3, glucose 20, NaHCO3 26, CaCl2 2.5 and bubbled with carbogen gas. Recordings were performed in a potassium gluconate based internal for both voltage clamp and current clamp recordings (K-gluconate 135 mM, NaCl 8 mM, HEPES 10 mM, Na3GTP 0.3 mM, MgATP 4 mM, EGTA 0.3 mM). Cells were visualized with a Sutter SOM microscope at 40x under IR illumination, and optogenetic stimulation was conducted through the microscope’s fluorescence path using a thorlab LED with a center wavelength of 565 nm. Data was acquired and filtered at 2.2 kHz using a Multiclamp 700B Amplifier (Axon Instruments) and digitized at 20 kHz (National Instruments Digitdata 1550B). All data were acquired using WinWCP and analyzed with custom python scripts.

### Organoid culture

Embryoid body formation and retinal organoid differentiation was performed as previously described^99^ with minor modifications. Briefly, iPS15 stem cells (ALSTEM) were dissociated to a single cell suspension with ACCUTASE, centrifuged at 300 RCF for 5 minutes, and resuspended in mTeSR+ with 10 µM Y-27632. On day 0, cells were seeded at a concentration of 10,000 cells/well in round bottom, ultra low attachment 96 well plates (Costar) at a volume of 100 µl per well. Plates were centrifuged at 100 RCF for 3 minutes and incubated at 37°C, 5% CO_2_ overnight. On day 1, 50 µl of media was removed from each well, and 150 µl of 1:1 mTeSR+ to neural induction media [NIM, composed of DMEM/F-12 with 1% v/v N2 supplement, 1% v/v MEM non-essential amino acids (Sigma) and 0.5% w/v heparin (Sigma)] was added to each well. 66.67% of the media was changed every other day with NIM until day 9. On day 9, organoids were transferred to a 1% Geltrex coated 6 well culture plate, plating 16 organoids per well, in 3 mL of NIM. 66.67% of the media was changed every other day with NIM until day 17. On day 17, 66.67% of the media was changed with organoid retinal induction medium (oRIM), composed of 1:1 DMEM basel media to DMEM/F-12, 2% B27 supplement without vitamin A, 1% Glutamax, 1% Penicillin/Streptamycin, 1% Normacin. 66.67% of the media was changed every other day with oRIM until day 23. On day 23, optic vesicle structures were scraped with a cell scraper and transferred to low-adhesion solution (STEMCELL Technologies) coated 6 well plates in oRIM supplemented with 2.6 nM (20 ng/mL) IGF1 (R&D Systems). On day 24, 66.67% of the media was changed with oRIM supplemented with 20 ng/mL IGF1. 50% of the media was changed with oRIM supplemented with 20 ng/mL IGF1 every other day until day 36. Starting on day 36, 50% of the media was changed with oRIM supplemented with 2.6 nM IGF1, 0.1 mM Taurine (Sigma), and 10% FBS every other day until day 66. Starting on day 66, 50% of media was changed with oRIM supplemented with 2.6 nM IGF1, 0.1 mM Taurine, 1 µM 9-cis retinal (Sigma), and 10% FBS, every other day until day 94. On day 94, 50% of the media was changed with oRIM supplemented with 2.6 nM IGF1, 0.1 mM Taurine, 0.5 µM 9-cis retinal (Sigma), and 10% FBS. Starting on day 96, 50% of the media was changed with organoid retinal induction maturation media (oRIM2), composed of 1:1 DMEM basel media to DMEM/F-12, 1% (v/v) N2 supplement, 1% (v/v) Glutamax, 1% (v/v) MEM non essential amino acids, 1% Penicillin/Streptamycin (v/v), 1% (v/v) Normacin, and supplemented with 2.6 nM IGF1, 0.1 mM Taurine, 0.5 µM 9-cis retinal (Sigma), and 10% FBS, every other day until organoids were harvested for downstream applications.

Organoids were transferred to individual wells (1 organoid per well) of a round-bottom, ultra-low attachment 96 well plates in 200 µL of media, between days 40-60. These were individually exposed to virus infection at a concentration of 5e10 vg/organoid. 50% of the media was changed after 2 days, and the regular organoid differentiation steps were performed as indicated above. Tips were changed between wells to avoid cross contamination between viruses and organoids. 4 weeks after virus exposure, the organoids were either fixed for histology or dissociated for single cell sequencing analysis.

### Microscopy

Fluorescent images were captured using a ZEISS LSM 980 confocal microscope with the MPLX Airyscan 2.0 super resolution detector. Micrographs were captured using the Plan-Apochromat 10x/0.45 air and 20x/0.8 air objectives (Carl Zeiss Microscopy, Jena, Germany). Laser lines used for fluorescent excitation included 405, 488, 561, 594 and 639 nm, with corresponding emission bandpass filters of 380-548nm, 495-550nm, 530-582nm, 673-627nm and 607-750nm, respectively. Tilescan images of larger areas of tissue were taken with a 10% overlap with pixel dimensions of 0.487µm(x) x 0.487µm(y) x 1µm (z). Single tile images had pixel dimensions of 0.122µm(x) x 0.122µm(y) x 0.5µm (z). ZEN Blue 3.5 software was used to perform Airyscan processing (auto-filter and standard strength settings), stitching of tilescans and maximum intensity projections of Z-stacks.

### Sequencing

#### Rabbit Retina dissociation

Rabbit retinas were dissected and cut into small pieces prior to dissociation. Dissociation was done using the Papain Dissociation System (Worthington Biochem), with slight modification. Briefly, retinal tissue was placed in papain/DNAse solution as per manufacturer’s instructions, and incubated on a thermal shaker at 37°C at 300 RCF for 60 minutes. The dissociated tissue was gently pipetted 10 times with a 10 mL serological pipette, and the supernatant was strained with a 40 µm cell strainer into a centrifuge tube. A discontinuous density gradient was prepared according to the manufacturer’s instructions, and the supernatant was discarded and the pellet was resuspended in PBS containing 1% BSA. The cells were counted and diluted to a concentration of 1000 cells / µL.

#### Single cell library preparation and sequencing

Single cell libraries were prepared using the 10x Chromium Next GEM Single Cell 3’ Kit, v3.1 and associated components. Quality control was performed on completed libraries using the Bioanalyzer High Sensitivity DNA Analysis kit (Agilent). Libraries were pooled and loaded onto an Illumina NextSeq1000 P2 100 cycle flow cell. Sequencing was performed in a paired end and dual indexes format, using the Illumina DRAGEN FASTQ Generation - 3.8.4 workflow with the Single Cell RNA Library Kit 1 and Single Cell RNA Index-Adapters 1-B kit.

### Sequencing analysis

A custom genome assembly was generated by putting together rabbit reference genome (2.0.107) with the sequence of the virally encoded transcript (i.e., ChRmine:mScarlet:Kv2.1-tag:WPRE:BGHpA) using Cell Ranger (7.0.1). The sequencing reads rabbit retina, previously injected with the AAV, were then aligned to this custom genome through Cloud Analysis platform from 10x Genomics. The dataset underwent normalization, variable feature selection, and scaling using *NormalizeData*, *FindVariableFeature*, and *ScaleData* functions from Seurat. Putative doublets were identified and eliminated using the DoubletFinder package^101^. The identities of the clusters were identified using a set of marker genes.

### Animals

C57/B6J and NCG male and female mice obtained from Charles River were used in this study. SPF New Zealand White female rabbits obtained from Western Oregon Rabbit company and Envigo weighing between 2-4kg were used in this study. Animals were maintained on a 12 hour light-dark cycle. All animal procedures were carried out with the approval of Science Corporation’s institutional animal care and use committee.

### ECoG Implantation Surgery

Implantation of surface grids (E32-1000-30-200, NeuroNexus, Ann Arbor, MI) was carried out under aseptic conditions. One hour prior to surgery, animals were given a dose of dexamethasone (2 mg/kg) to curtail brain swelling. Animals were anesthetized with a cocktail of ketamine (20 mg/kg), xylazine (3 mg/kg), and acepromazine (2 mg/kg) and placed in a stereotaxic frame (Model 1240, Kopf Instruments, Tujunga, CA) fitted with a custom 3D printed nose cone and maintained with isoflurane inhalation (1-5% in O_2_) throughout. Animals rested on a heating pad at 35-37 C° for the entirety of the surgery. A midline incision from the center of the eyes to the occipital suture was made and the cranium exposed through blunt dissection of the underlying tissue. We first secured a custom head holder (titanium) to the nasal bone with cortical screws for head fixation during electrophysiological procedures. Next, a craniotomy (bregma −10 mm to lambda +2 mm, 2 to 10 mm from the midline) exposed visual cortex. Following durotomy, the grid was placed and the dura replaced. ECoG responses elicited by gross visual stimulation of the contralateral eye confirmed grid placement. An artificial dura (DuraGen, Integra Life Sciences, Princeton, NJ) was placed in the craniotomy, and a custom chamber was secured to the cranium via acrylic resin. The chamber was then filled with silicone elastomer (Kwik Sil, World Precision Instruments) and closed with a cap and silicone spacer. The wound margin was sutured and the animal removed from the stereotax. Animals were given subcutaneous long-acting analgesics (buprenorphine slow release, 0.12 mg/kg) and antibiotics (enrofloxacin, 5 mg/kg) and returned to their home cages once sternal, and received analgesics (meloxicam, 5 mg/kg) once daily for 3 days post surgery. Sutures were removed 10 days post-surgery once wound margins were healed.

### Intravitreal injections

*Mouse:* Intravitreal injections of AAV in mice were performed under isoflurane anesthesia (age, 3-4 months old). The procedures typically lasted about 5 minutes. Prior to injection, we applied a drop of proparacaine hydrochloride ophthalmic solution to the eye (0.5%). AAV solution was injected intravitreally using a beveled glass micropipette loaded on a microinjector (Nanoject II, Drummond; speed, 70 nl/s). The micropipettes were pulled such that the tip diameter was between 50 and 100 microns. The animals were allowed to recover in their home cage for at least 2 weeks prior to any experiment.

#### Rabbit

Under general anesthesia eye drops were applied to dilate the iris (atropine 1%, tropicamide 1% and phenylephrine 10%) and promote topical anesthetic (proparacaine 0.5%). A surgical microscope (Zeiss OPMI Visu 160 on a S7 Stand) was used to fully insert a 30g 5 / 8 length needle through the sclera (1.5mm from the limbus) and solutions (sodium iodate, AAV virus, synaptic blockers) were injected into the vitreous cavity with the needle end aimed towards the visual streak. After injection the needle was slowly extracted from the sclera. Intravitreal injections were either virus (1e11 vg /eye), 0.4 mg sodium iodate, or a cocktail of synaptic blockers in volumes less than 150 µL. For synaptic blockers, concentrations were assumed to be diluted 12x based on the putative volume of the rabbit eye (1.76 mL based on axial diameter of 15 mm measured by A-scan (DGH 6000, ScanmateA)). The working concentration in the eye of synaptic blockers was in mM (AP5: 0.05, AP4: 2, GYKI 0.05, NBQX: 0.01, UBP310: 0.01, PDA (cis-2,3-piperidine dicarboxylic acid): 10).

### Ophthalmic procedures

Intraocular pressure (IOP) was measured in rabbits during general anesthesia using tonometry (tonopen, Reichert) and central corneal thickness (CCT) using pachymetry (DGH Pachmate 2). For both procedures rabbits were either in a lateral decubitus position or prone position.

### Visual stimulation

To test retinal function, we performed electroretinograms (ERG; RETIport 3S, An-Vision, West Jordan, UT) using small animal Jet electrodes (Fabrinal Eye Care, La Chaux-De-Fonds, Switzerland). We performed examinations under both photopic and scotopic conditions. For photopic vision assessment, animals were sedated in the operating room after a minimum of 10 minutes light adaptation and tested with a standard flash (0dB) to access combined rods and cones, and a flicker flash (0dB, 28Hz) to isolate cone response, with a background light intensity of 25 cds/m^2^. For examination of scotopic vision which isolates rod vision, animals were sedated in the housing room in the last 30 minutes of their normal dark cycle (12 hours dark adaption) and two types of light intensity were used; a small light intensity (−20bB), followed by a standard flash (0dB) measuring both maximal and oscillatory potentials.

We employed a battery of visual stimulations to assess visual cortex local field activity under several longitudinal conditions. First, to assess baseline V1 function, anesthetized animals were stationed sternally with a 24 inch 240 Hz LCD monitor positioned approximately 10 cm from the eye. In order to account for the focal length of the rabbit eye, all stimuli were spherically corrected to ensure equal weighting in the neural response. Receptive field mapping consisted of a white circle presented pseudorandomly in visual space along a 9 x 7 grid spanning roughly 80 horizontal and 60 vertical degrees of the visual field. Each stimulus was presented for 200 ms and spanned 10 degrees of visual angle, followed by a variable interstimulus interval ranging from 300 to 500 ms. We performed stimulation at each location on the grid 25 times (1575 trials total). Next, we presented full field moving gabor filters with a spatial frequency of 0.01 degrees whose orientation and hence direction of motion was pseudorandomly selected from 0 to 150 degrees in increments of 30 degrees. Each gabor was repeated 30 times. We also presented full field white light stimuli by varying the grayscale intensity of a full field box stimulus ranging from 0 (black; no change from screen background) to 1 (white) with 11 levels, repeating each stimuli 30 times. This range corresponded to luminance values of 8.6 to 641 lumens. All visual stimuli were presented using the python psychophysics platform PsychoPy^103^ using the coder interface.

### Optogenetic stimulation

In order to test virus expression in the rabbit, we first performed full field visual stimulation using a 530 nm LED (M530L4, ThorLabs, Inc., Newton, NJ) passed through a collimating lens with a focal length of 30 mm. Given that the axial length of the rabbit eye is approximately 15 mm, we positioned the LED 15 mm from the eye. Stimulation consisted of short-duration 50 ms pulses, ranging from 2.2 to 58.7 mW/cm^2^ in 11 uniform increments. Forty to 50 repetitions of each stimulus power were performed in a pseudorandomized order with an interstimulus interval of 300 to 500 ms. Stimulus powers were determined using PsychoPy and an Arduino Due which provide pulse width modulation (PWM) input to the LED.

Two rabbits underwent postoperative electrophysiological testing following FlexLED implantation, prior to recovery from anesthesia. The stimuli consisted of turning on all active pixels on the FlexLED device for either 200 or 500 ms.

### Histology

#### Mouse Flat Mount Retina Histology

Experiments were carried out on wild type C57/B6J mice at 1, 2, and 4 weeks post-injection. Animals were euthanized by elevated carbon dioxide and cervical dislocation. Eyes were quickly enucleated and the retinas were dissected free of the vitreous and sclera in carboxygenated Ames Medium (A1372-25 US Biologica). Four cuts were made in each retina to allow the samples to remain flat and mounted onto 0.5 x 0.5 cm hydrophilic filter paper (MF-millipore 0.45 µm MCE Membrane, 47 mm). Samples were then fixed in 4% paraformaldehyde (PFA) for 1 hour, then washed three times in PBS for 5 minutes. Whole mounted retinas were then blocked in a solution containing 10% donkey serum (DS), 0.1M Glycine, 0.3% Triton X-100 in PBS for 1 hour. Primary antibodies were diluted in 1% DS in PBS overnight. Samples were washed in PBS three times for 5 minutes, then incubated for 2 hours at room temperature with secondary antibodies diluted in PBS. Samples were washed in PBS three times for 5 minutes, then 4’,6-Diamidino-2-Phenylindole, Dihydrochloride (DAPI) (D1306 Thermofisher Scientific) (1:1000) was added to all the samples to stain cell nuclei and incubated at room temperature for 5 minutes. All samples were washed again with PBS for 5 minutes. Retinas were then transferred and mounted onto glass slides with a drop of ProLong™ Gold Antifade Mountant (P10144, Thermofisher scientific), and then covered with a glass coverslip.

#### Rabbit Whole Mount

Experiments were carried out on female SPF New Zealand White Rabbits with post intravitreal injections at 4 week time points. Animals were euthanized with Pentobarbital overdose (200mg/kg; 2.5ml) IV. Eyes were quickly enucleated and 4 cuts were made in the sclera to allow opening of the eyecup. Dissections were performed in carboxygenated Ames Medium (A1372-25 US Biologica). The retina was separated from the choroid and the sclera using a fine paintbrush to prevent retina tissue damage. Equally sized samples were taken from four regions of the retina using 3mm biopsy punches. Biopsy punched retina samples were mounted onto 0.5×0.5cm hydrophilic filter paper (MF-millipore 0.45µm MCE Membrane, 47mm). Such samples were then fixed in 4% paraformaldehyde (PFA) for 1 hour. Samples were washed three times in PBS for 5 minutes. Whole mount retinas were then blocked in a solution containing 10% donkey serum (DS), 0.1M Glycine, 0.3% Triton X-100 in PBS for 1 hour. Primary antibodies were diluted in 1% DS in PBS overnight. Samples were washed in PBS three times for 5 minutes, then incubated overnight at room temperature with secondary antibodies diluted in PBS. Samples were washed in PBS three times for 5 minutes, then 4’,6-Diamidino-2-Phenylindole, Dihydrochloride (DAPI) (D1306 Thermofisher Scientific) (1:1000) was added to all the samples to stain cell nuclei and incubated at room temperature for 5 minutes. All samples were washed again with PBS for 5 minutes. Retinas were then transferred and mounted onto glass slides with a drop of ProLong™ Gold Antifade Mountant (P10144, Thermofisher scientific), and then covered with a glass coverslip.

#### Rabbit Cryosections

Experiments were carried out on female SPF New Zealand White Rabbits with post intravitreal injections at 4 week time points. Animals were euthanized with Pentobarbital overdose (200mg/kg; 2.5ml) IV. Eyes were quickly enucleated in carboxygenated Ames Medium (A1372-25 US Biologica). Samples were fixed in 4% paraformaldehyde (PFA) for 1 hour. Samples were washed three times in PBS for 5 minutes. Samples were cryoprotected in incremented concentrations of sucrose solutions. Samples were placed in a 10% sucrose solution for 1hr and 4°C, proceeded by placing the samples in a 20% sucrose for 1hr at 4°C and then finally placed in a 30% sucrose solution overnight. Samples were next embedded in OCT in plastic cryomolds and snap frozen in ⅔ isopentane and ⅓ liquid nitrogen mixture. Samples were left in the −20°C freezer overnight. Using a Leica cryostat, 50µm cryosection retinal slices were made and mounted onto Fisherbrand™ Superfrost™ Plus Microscope Slides (Fisher Scientific Cat No. 12-550-15). Samples on slides were then blocked in a solution containing 10% donkey serum (DS), 0.1M Glycine, 0.3% Triton X-100 in PBS for 1 hour. Primary antibodies were diluted in 1% DS in PBS at room temperature overnight. Primary antibodies were diluted in 1% DS in PBS overnight. Samples were washed in PBS three times for 5 minutes, then incubated for 2 hours at room temperature with secondary antibodies diluted in PBS. Samples were washed in PBS three times for 5 minutes, then 4’,6-Diamidino-2-Phenylindole, Dihydrochloride (DAPI) (D1306 Thermofisher Scientific) (1:1000) was added to all the samples to stain cell nuclei and incubated at room temperature for 5 minutes. All samples were washed again with PBS for 5 minutes.

#### Organoid Cryosections

Organoids were fixed in 4% PFA for 2 hours on a shaker at room temperature. Samples were washed three times in PBS with 0.1M Glycine for 5 minutes. Samples were placed in 30% sucrose solution overnight at 4°C. Samples were then embedded in OCT in plastic cryomolds and stored at −20°C for at least 24 hours. Using a Leica cryostat, 13 µm cryosection organoid slices were made and mounted onto Fisherbrand™ Superfrost™ Plus Microscope Slides. Slides were dipped in purified water to wash away residual OCT. Samples were then blocked in a solution containing 10% DS, 0.5% Triton X-100, and 0.5% BSA in PBS for 1 hour. Primary antibodies were diluted in 1% DS, 0.5% Triton X-100, and 0.5% BSA in PBS and added to the samples at 4°C overnight. Samples were washed three times in PBS for 5 minutes. Secondary antibodies were diluted in 1% DS, 0.5% Triton X-100, and 0.5% BSA in PBS and added to the samples at 4°C overnight or for 2 hours at room temperature. Samples were then washed three times in PBS for 5 minutes, and stained with DAPI (1:1000) for 10 minutes at room temperature. Samples were washed three times with PBS for 5 minutes, and mounted with coverslips using ProLong™ Gold Antifade mounting media.

**Table 2:** Primary Antibody List

| **Antigen** | **Host** | **Target Cell** | **Dilution** | **Source** | **Cat No.** |
| --- | --- | --- | --- | --- | --- |
| Brn3a | Rabbit | Retinal Ganglion cells | 1:100 | Abcam | ab245230 |
| RBPMS | Guinea Pig | Retinal Ganglion cells | 1:100 | Phosphosolutions | 1832-RBP MS |
| HuC/HuD | Mouse | Retinal Ganglion Cells | 1:200 | Thermofisher | A-21271 |
| GFAP | Chicken | Glia | 1:1000 | Abcam | ab7260 |
| MAP2 | Mouse | Neurons | 1:1000 | Millipore Sigma | M1406 |
| BLIMP1 | Rat | Photoreceptors | 1:100 | Santa Cruz Technology | sc-130917 |
| NRL | Goat | Rod Nuclei | 1:500 | R&D Systems | AF2945 |
| CD3 | Rabbit | T cells | 1:1000 | Biocare Medical | CP215A |
| CD20 | Mouse | B cells | 1:400 | DAKO | M0755 |
| IBA1 | Goat | Microglia/macro phage | 1:1000 | Abcam | ab5076 |
| Recoverin | Mouse | Photoreceptors | 1:1000 | Abcam | ab31928 |
| Recoverin | Rabbit | Photoreceptors | 1:1000 | Millipore | AB5585-I |

**Table 3:** Secondary Antibody List

| Species | Target | Fluorochrome | Dilution | Source | Cat. No |
| --- | --- | --- | --- | --- | --- |
| Recombinant | Anti-Rabbit | Alexa Fluor 647 | 1:500 | Abcam | ab300158 |
| Donkey | Anti-Rabbit | Alexa Fluor 594 | 1:500 | Abcam | ab150064 |
| Goat | Anti-Guinea Pig | Alexa Fluor 488 | 1:500 | Abcam | ab150185 |
| Donkey | Anti-Mouse | Alexa Fluor 488 | 1:500 | Abcam | ab150105 |
| Donkey | Anti-Chicken | Alexa Fluor 488 | 1:500 | Biotium | 20166 |
| Goat | Anti-Rat | Alexa Fluor 546 | 1:500 | Thermofisher | A-11081 |
| Donkey | Anti-Mouse | Alexa Fluor 647 | 1:200 | Abcam | ab150107 |

### FlexLED Implantation

Animals were anesthetized with a cocktail of ketamine (20 mg/kg) and xylazine (3 mg/kg), dilating eye drops (atropine 1%, tropicamide 1%, phenylephrine 10%) and topical anesthetic drops (proparacaine 0.5%) were applied to the surgical eye, and hydro gel (genteal) applied to both eyes. The hind leg was shaved and a catheter placed for IV fluids and the patient transported to the operating room. The patient was positioned in lateral decubitus position and general anesthesia maintained via isoflurane delivery through a nose cone or intubation. The operative eye was prepped (betadine) and draped. Using sterile technique, eyelids were retracted using five 1mm blunt retractors (fine science tools). A surgical microscope (Zeiss OPMI Visu 160 on a S7 Stand) was used for visualization. Wescott scissors were used to create a superior 180° conjunctival peritomy. A paracentesis was created using a 1.1 mm blade and epinephrine was injected into the anterior chamber to maintain iris dilation. Two 25g trocars are placed 1.25 mm from the back of the limbus in the superotemporal and superonasal quadrants for intraocular access. Viscoelastic was injected into the anterior chamber and a temporal clear corneal incision was fashioned with a 2.75 mm metal keratome superiorly. An anterior capsulotomy was created with a cystotome needle and lehner-utrata capsulorhexis forceps using a can opener technique. The lens was then removed using the phacoemulsification handpiece. The main clear corneal incision was closed with a 10-0 nylon suture in figure-eight fashion and the knot was buried. An anterior chamber maintainer is inserted through a side port incision into the cornea to irrigate the globe with sterile balanced salt solution (BSS) and epinephrine (500:1). A 25g vitrector and light pipe were introduced into the eye through the trocars and the lens capsule was removed. Next, a core vitrectomy was performed with assistance of the Oculus Biom Ready system. Dilute Triesence was then administered and used to stain the vitreous, allowing careful removal of remaining vitreous.

A 18g cautery is used to pretreat the sclerotomy site, irrigation is stopped and BSS replaced with viscoelastic (Amvis plus). A mattress suture (7-0 vicryl) was pre-placed around the sclerotomy site using partial thickness scleral bites. A 5 mm scleral incision is performed in the superotemporal pars plana 1.5mm from the limbus using a super shape blade. The FlexLED supported by the rigid carrier device was gripped with tying forceps at the carrier grip points and the intraocular portion of the device was carefully inserted through the sclerotomy. Visco elastic was re-applied to the eye to maintain pressure. The anterior aspect of the extraocular device was secured to the sclera with non dissolving 5-0 mersilene suture through the anterior and 7-0 vicryl suture through the posterior anchoring loops. The carrier device was released from the device by cutting pre-placed sutures in the anterior anchoring loops. Tying forceps were used to grip the device while the carrier ripcord was pulled to release the intraocular portion of the device. The pre-placed mattress suture was then tied in 3-1-1 fashion to seal the sclerotomy. Visco elastic was re-applied to the eye to maintain pressure. A light pipe and intraocular forceps were introduced through the trocars and used to appropriately position the intraocular FlexLED array on the retina, with care to contact only the pre-designated grab site on the device. A small peritomy was made inferiorly opposite the device insertion site and 18g cautery was used to pretreat this area before a super sharp was used to make the sclerotomy ∼1.5 mm, 1.25 mm from the limbus. A pair of tacking forceps were loaded with a retinal tack (length: 2.8 mm, maximum width: 0.8 mm) and introduced into the eye through the inferior incision. The intraocular device was tacked through the tack hole to the retina. BSS infusion was resumed and a vitrector was used to remove viscoelastic from the globe. All sclerotomies were closed with 7-0 vicryl suture and the eye was made watertight. The FlexLED power cable was tucked next to the device, the conjunctiva was draped over the device and sutured to the limbus with interrupted 7-0 vicryl sutures. Subconjunctival injections of triamcinolone, moxifloxacine and lidocaine were administered. The retractors and drapes were removed. The subject was monitored for recovery for 1 hour and transported to the housing facility in stable condition. Animals received dorzolamicide and brimonidine drops once a day for 3 days after surgery, NeoPolyDex five times a day for three weeks post surgery, and phenylephrine and tropicamide twice a day for three weeks post surgery.

### *In vivo* electrophysiology and analysis

We performed ECoG recording during the anesthetized procedures described above. ECoG signals were notch filtered at 60 Hz and digitized at 4-5 kHz via a 32-channel recording headstage and a 1024-channel RHD recording controller (Intan Technologies, Los Angeles, CA) and streamed to disk on a Linux-based PC. Stimulus timings were digitized via Arduino input into either auxiliary digital or analog ports on the recording controller. In order to capture event times streamed from the FlexLED, signals were sampled at 30 kHz. In most rabbits, we performed electrophysiological recordings at a minimum of two timepoints: post ECoG implant, and at least 10 days following sodium iodate injection. The former session allowed us to estable baseline responses to full field visual stimulation (gratings, luminance), to perform a map of visual receptive fields, and to test endogenous responses to 530 nm LED stimulation. The latter session, which followed selective destruction of photoreceptors, allowed us to confirm the absence of white-light visual stimuli responses, and to titrate the cortical responses evoked from activation of opsin-expressing RGCs. Two rabbits underwent synaptic blockade to provide a further control over cortical responses arising from any spared photoreceptors following SI injection. ECoG recordings were taken in the two rabbits implanted with the FlexLED device.

Data analyses were carried out in Python. To preprocess, we wideband filtered between 3 and 500 Hz using a bidirectional 5th order Butterworth filter after notch filtering the raw data at 60 Hz and its harmonics (IIR notch filter) and downsampling to 1 kHz. We then performed two additional processing steps, depending on the analysis. Analysis of gamma (20-80 Hz in the awake rabbit^104^, which is predictive of multi-unit activity in the brain^105^, was carried out by band-filtering and taking the absolute magnitude of the envelope using the Hilbert transform (Hilbert power). We analyzed wideband signals by decomposing into time-frequency components using Morlet wavelets^106^ with 6 cycles from 10 to 300 Hz in steps of 4 Hz across time windows of interest at a resolution of 1 ms. For visualization, we normalized time-frequency power by baseline activity. Processed data was put into trials and sorted based on experimental conditions.

To decode RF stimulus location from V1 activity, we used a support vector machine approach (SciKit-learn, LinearSVC) in an one-vs-rest configuration. This approach tests each condition against a pool of all other conditions. We extracted a number of measures from the time-frequency decomposition of each ECoG channel to generate a feature matrix. First, wideband LFP was windowed at 100 ms pre to 400 ms post stimulus onset and zeroed to baseline. We extracted the peak, the latency, and frequency of the response within the gamma band within the first 100 ms of stimulus presentation, along with a 10-sample time course of the mean gamma response within the same window. We also extracted the peak amplitude and latency in the LFP visual evoked response (< 50 Hz). Thus, for each ECoG channel there were 15 features extracted, resulting in a feature matrix of 480 features x 1575 trials. The data were tested 10 times each with a randomized seed, and compared to a bootstrapped analysis to determine chance levels for the data. To do this, we repeated the decoding analysis 50 times using shuffled features to predict actual positions. In order to quantify decoder performance, we computed the number of correct predictions for decoders predicting horizontal, vertical, or joint visual angles and compared to shuffled. To better understand errors in decoding, we tabulated the Euclidean distance between actual and predicted locations within the visual space (Figure 4d).

### Optical Modeling and Measurement

#### Aphakic Photochemical Hazard Function

Optical exposure limits to the retina were calculated according to ICNIRP Guidelines on Limits of Exposure to Incoherent Visible and Infrared Radiation^60^. Starting from action spectra for an aphakic (lens-less) eye, allowable epiretinal irradiances for blue-light photoretinopathy were computed for a single LED pixel and a full LED array for chronic exposure (t > 0.25 s). Of the two known mechanisms of retinal damage, blue-light photoretinopathy occurs at significantly lower levels of irradiance compared to thermally induced photoretinopathy, hence was identified as the dominant source of damage in the wavelengths between 300 and 700 nm. Biologically weighted effective radiance was calculated as a weighted average of the spectral distribution of the LED—a Gaussian distribution with a full-width at half-maximum value of 20 nm—and the aphakic retinal thermal hazard function.

#### Angular Emission Profile Measurement

A u-turn device with a single LED lit up was placed on the rotating axis of a servo motor. A power meter (Thor labs, PM100USB) positioned 5 cm away from the LED measured normally incident power across a 90-degree angle span with increments of 2 degrees. To confirm the angular emission profile in an immersion state, the single-pixel lit device was immersed in a 10 µM solution of AF568 (Lumiprobe) and imaged through a 610 nm long-pass optical filter (Chroma AT610lp). The LED was oriented in the horizontal direction to minimize the amount of direct LED emission collected by the objective.

#### Monte Carlo Optical Scattering in Tissue

A custom Monte Carlo multi-layer simulation environment traced individual photon paths as it interacted with material interfaces as well as scattering and absorbing agents^107^. 10^7^ photons were launched with an angular probability distribution function following an empirically measured angular emission profile. Layers in the illumination path from the LED emission layer to the retina were modeled as a series of parallel material planes defined by their characteristic thickness, refractive index, scattering coefficient, absorption coefficient, and scattering anisotropy (Table 4). The Henyey-Greenstein Phase Function commonly used to model optical scattering in neural tissue determined the scattering angle at each scattering event. The LED emission layer was modeled as 6-by-11 (hinge) and 15-by-19 (u-turn) µm^2^ rectangular Lambertian light sources encapsulated in 10 µm of glass. The retina was assumed to be in contact with the device.

**Table 4:** Layer characteristics

| Layer | Thickness ($\mu\text{m}$) | Refractive index | Scattering coefficient ($\text{cm}^{-1}$) | Absorption coefficient ($\text{cm}^{-1}$) | Scattering anisotropy |
| --- | --- | --- | --- | --- | --- |
| LED emission layer | 0 | - | - | - | - |
| Glass | 10 | 2 | 0.0 | 0.0 | 1.0 |
| Glass | 3 | 2 | 0.0 | 0.0 | 1.0 |
| Retina | 50 | 1.358 | 71.5 | 2.745 | 0.7 |

---

**Supplemental Figure 1:**
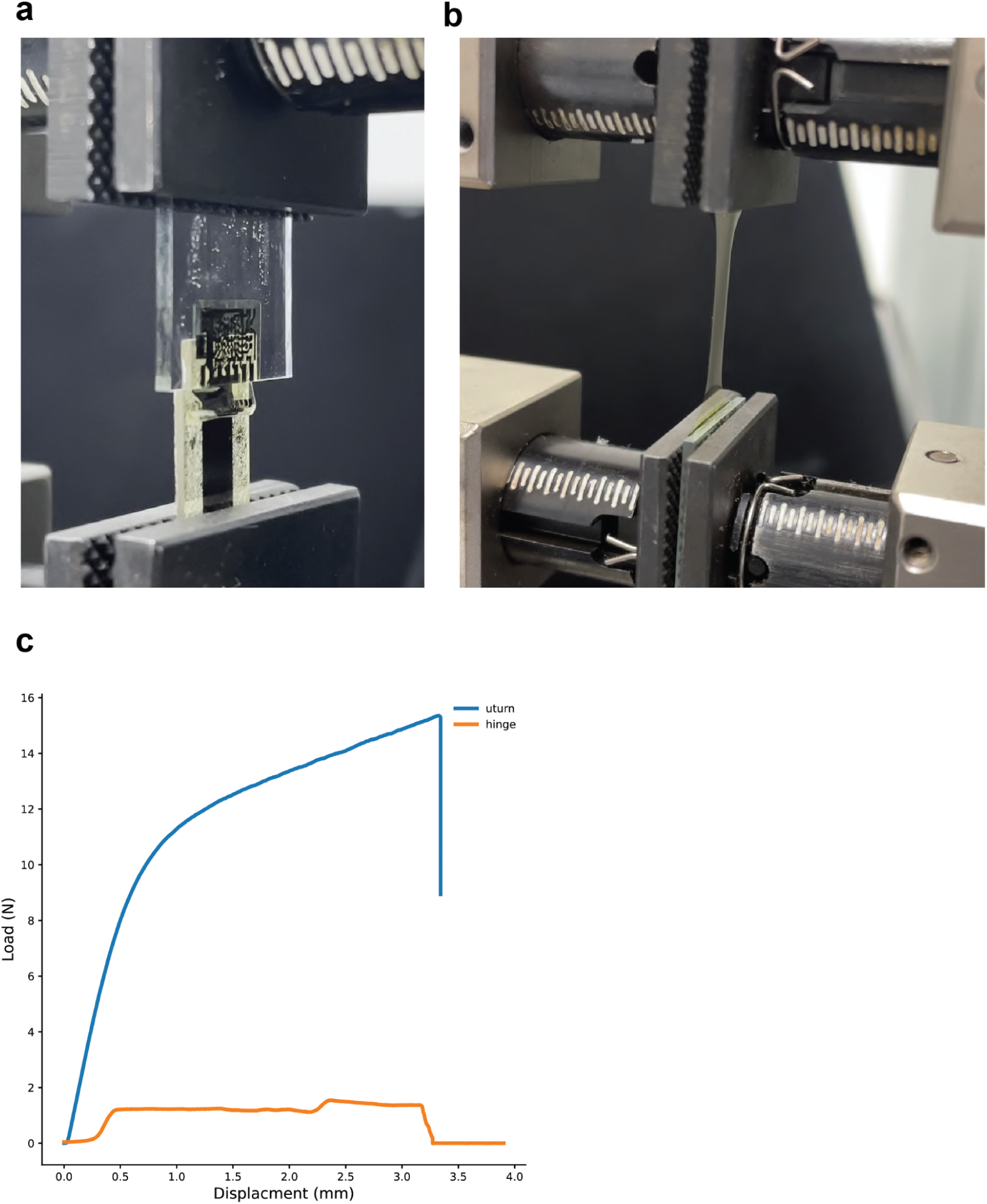
The u-turn device offers improved fracture load. a) Experimental peel test setup for characterizing adhesive force between the polyimide layers of the hinge design. b) Experimental tensile test setup for characterizing the yield point and fracture load of a polyimide structure of comparable cross-sectional geometry to the u-turn design. c) The u-turn design exhibits a 15x failure load over the hinge design.

**Supplemental Figure 2:**
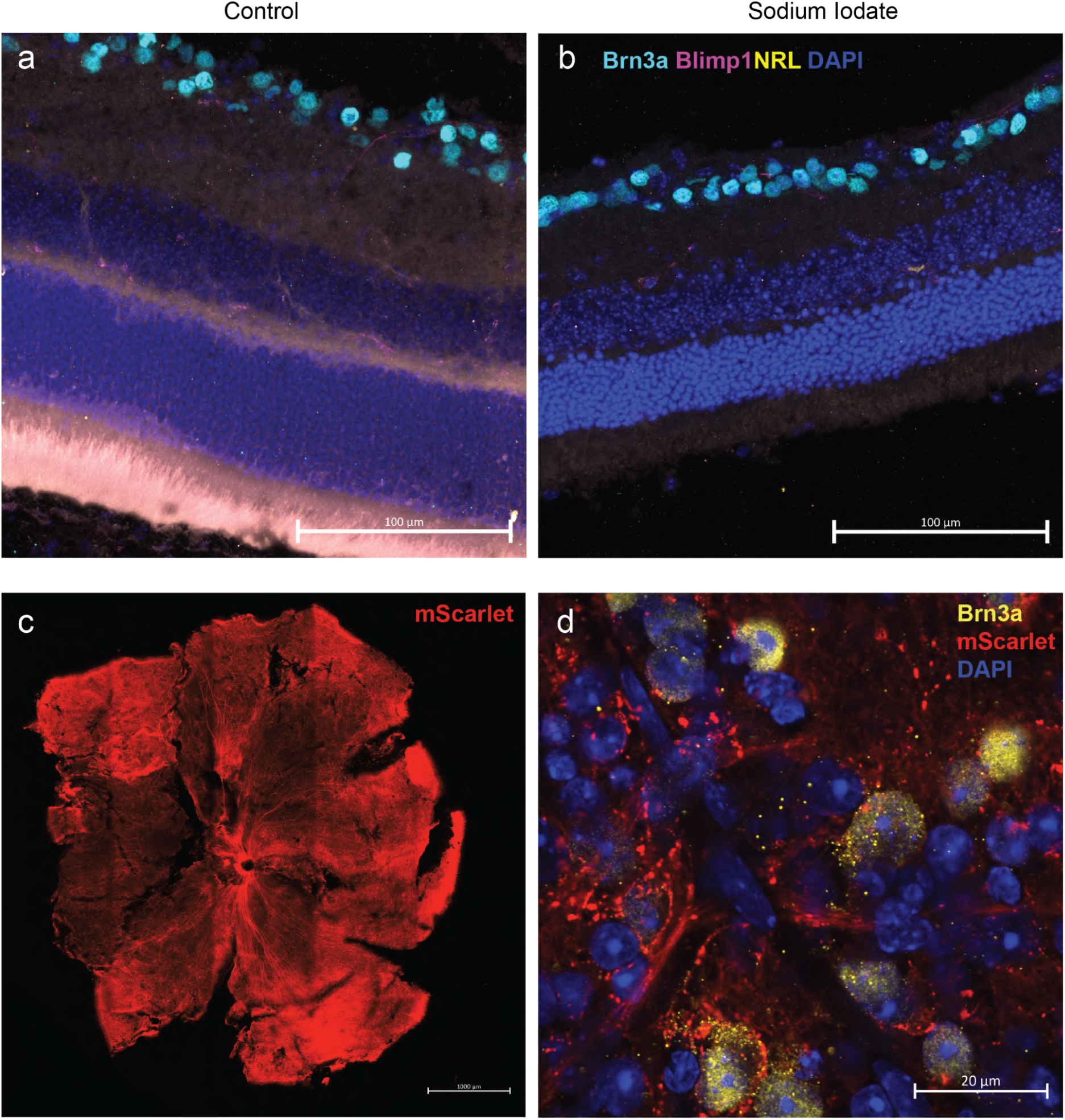
Validation of Sodium Iodate and Intravitreal injection of AAV vector in mice. a) Confocal microscope images from 20 µm thick cryosectioned retinas from a control mouse showing intact photoreceptors evident by expression of markers Blimp1 (magenta) and NRL (yellow) and RGCs (Brn3a, cyan). Scalebar 100 µm. b) As in (a), but a sodium iodate treated eye. c) Confocal microscope image from whole mount mouse retinas showing widespread viral transduction (mScarlet, scalebar 1000 µm) d) Confocal microscope image from whole mount mouse retinas showing widespread viral transduction (red) and colocalization with the RGC-marker Brn3a (yellow, scalebar 20 µm)

**Supplementary Figure 3:**
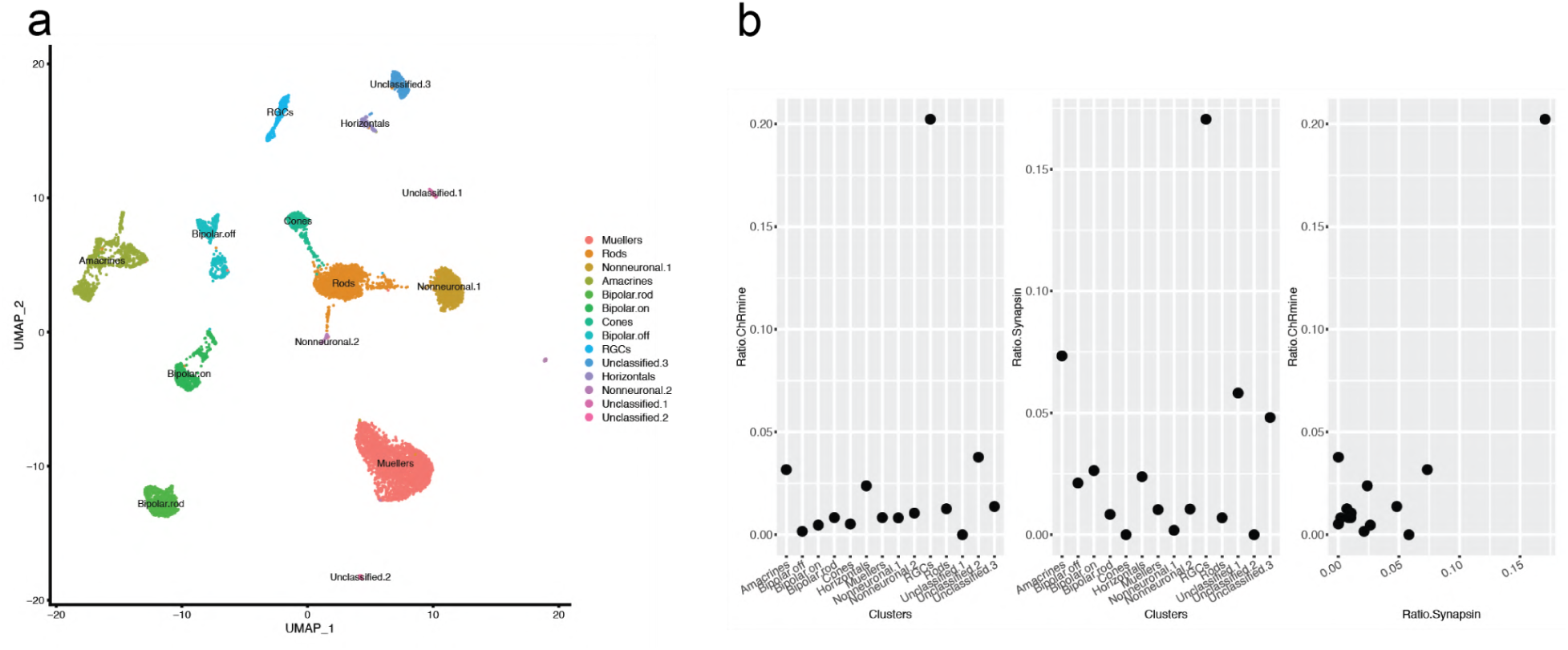
Single-cell sequencing from rabbit retina reveals selective expression of AAV-hSyn1-ChRmine-mScarlet-Kv2.1 in Retinal Ganglion Cells. a) Single cell seq data from rabbit retina (n=2 rabbits), dissociated for library prep about 6 weeks after Intravitreal injection of AAV-hSyn1-ChRmine-mScarlet-Kv2.1 (100 µl volume, 1.00E+11 GC/eye) b) RGC specific expression of ChRmine and Synapsin calculated as the ratio of the number of expressing cells by the total number of cells in each cluster.

**Supplemental Figure 4:**
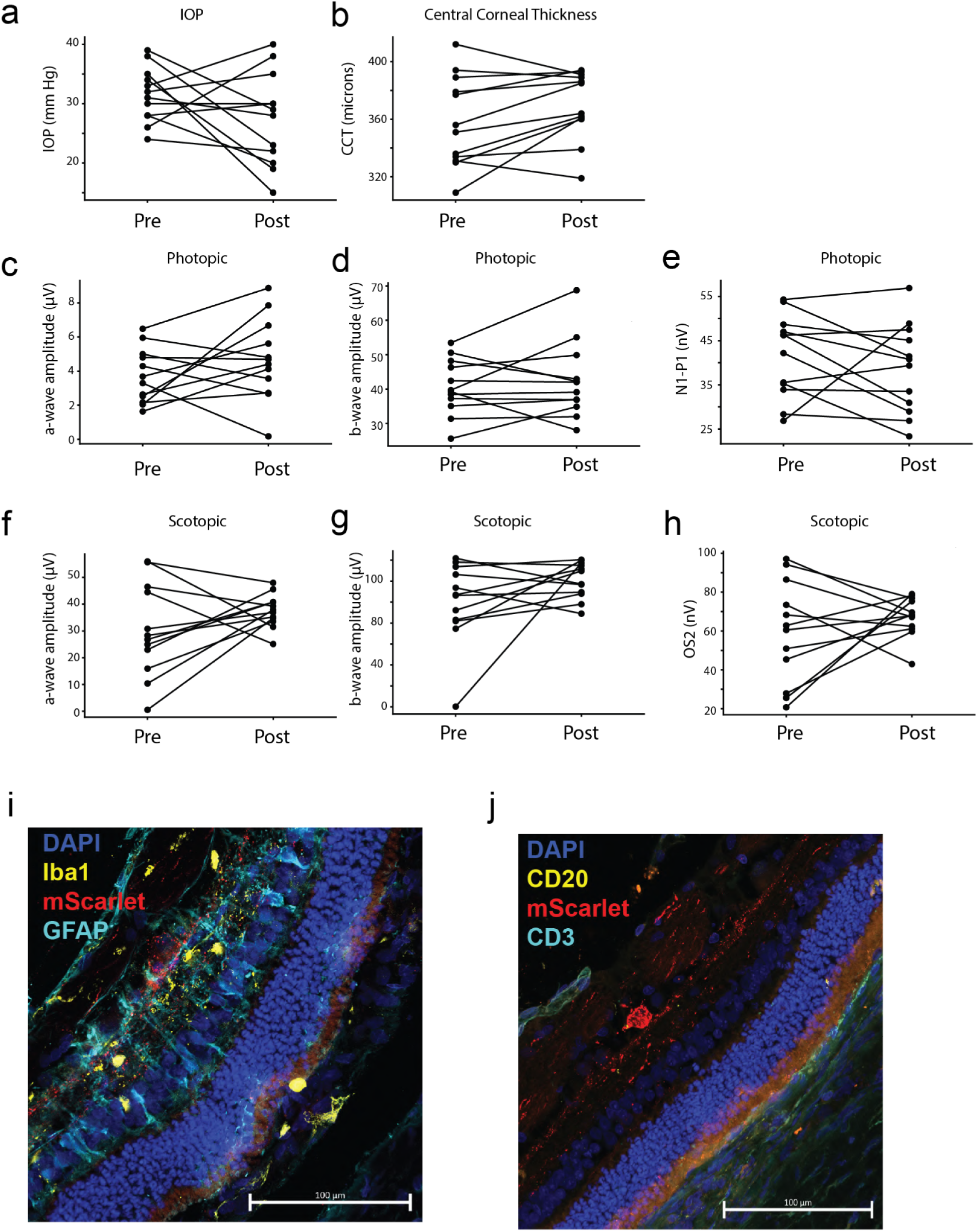
Intravitreal injection with AAV-hSyn1-ChRmine-mScarlet-Kv2.1 does not affect retinal function and does not cause an inflammatory response in rabbits Measurements taken before (pre) and 6 weeks after (post) intravitreal injections of AAV (N = 6 rabbits, 2 eyes / rabbit) a) Intraocular pressure (IOP) b) Central corneal thickness (CCT). c) Light adapted ERG response that were elicited by a bright (25 cd/m^2^) light stimulus with a standard flash (0 dB) measuring a-wave amplitude d) Light adapted b-wave amplitude e) N1P1 amplitude after a flicker flash (0 dB, 28 Hz) f) Dark adapted ERG response that was elicited by a standard flash (0dB) measuring the maximal potential a-wave amplitude g) Dark adapted b-wave amplitude h) Oscillatory potential OS2 amplitudes. i) Confocal microscope images from 20µm thick cryosectioned retinas from a rabbit after AAV transfection showing unremarkable, normal astrocyte (GFAP, cyan) and microglia (Iba1, yellow) response around opsin expressing cells. k) Markers for peripheral immune cells CD3 (cyan) and CD20 (yellow) showed no infiltration into the retinal space. Scale bar 100µm.

**Supplemental Figure 5:**
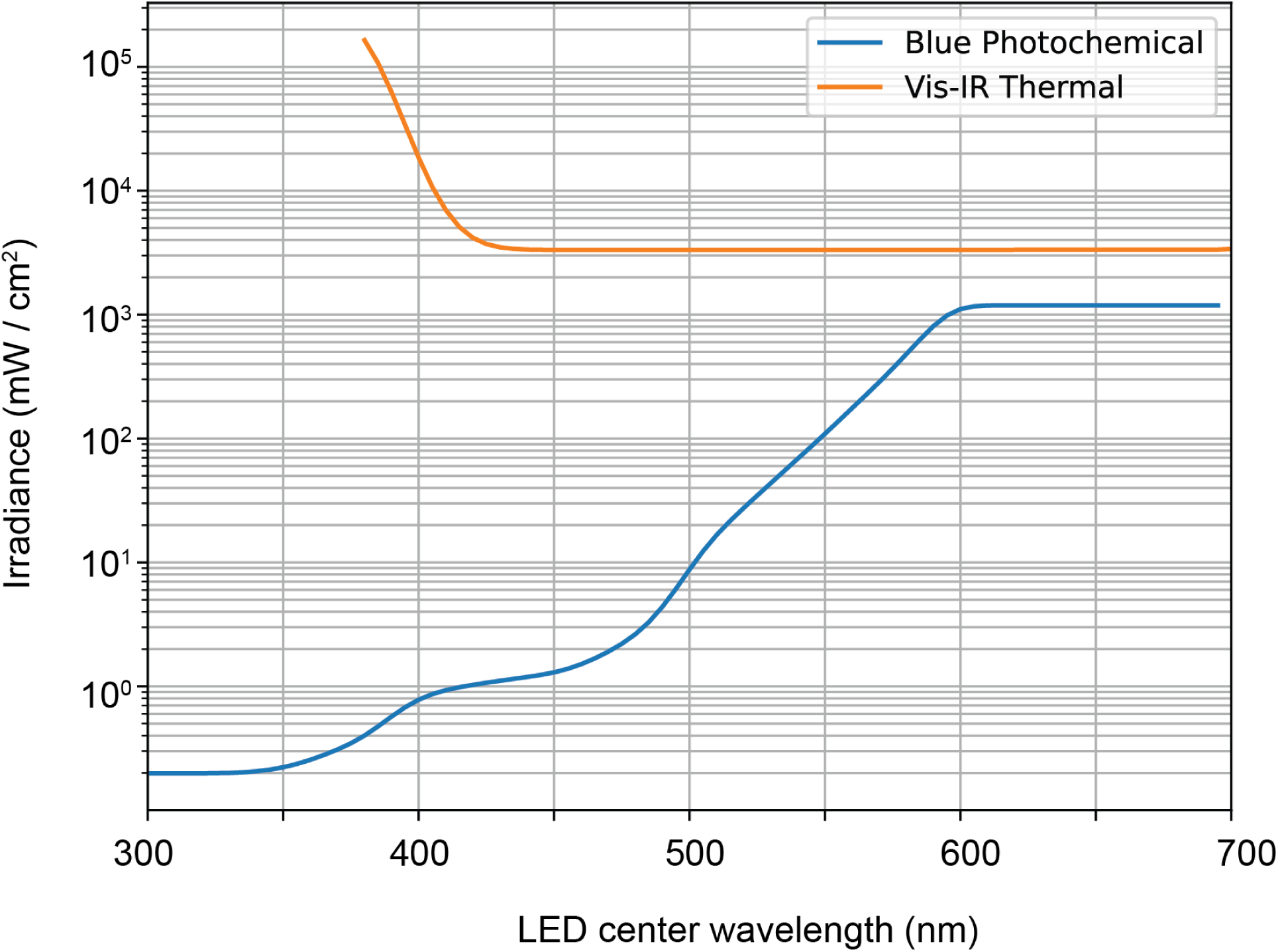
Optical safety limit of epi-retinal light source in compliance with International Commission on Non-Ionizing Radiation Protection Guidelines. Effective photochemical (blue) and thermal (orange) retinal exposure hazard functions are shown for visible light.

**Supplemental Figure 6.**
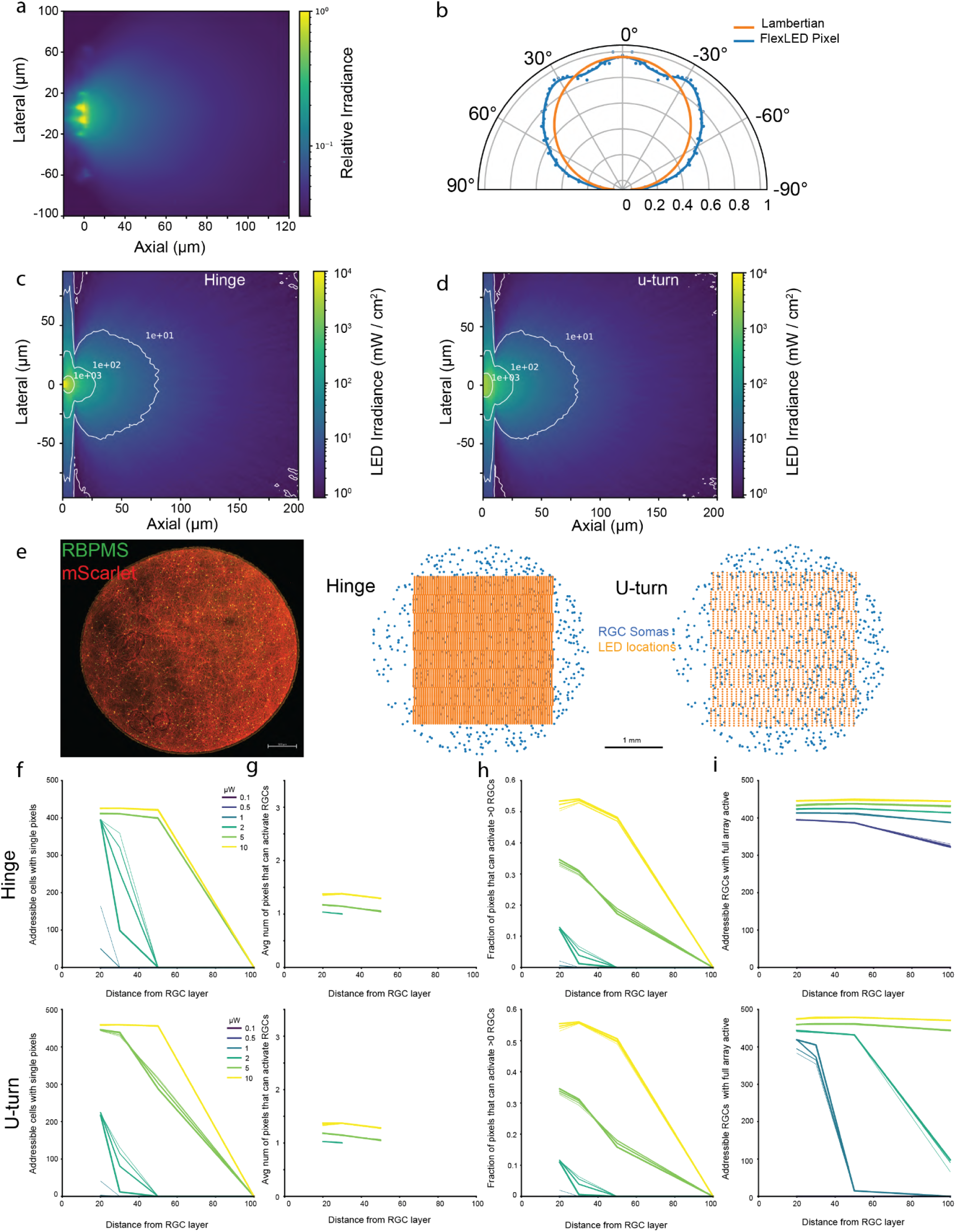
Modeling of FlexLED optogenetic stimulation from an empirically measured LED angular emission profile. a) Cross-sectional illumination profile of a single 15 x 19 µm u-turn device pixel immersed in a fluorescent solution b) Polar plot comparing the angular emission profile of a u-turn pixel (blue) to an ideal Lambertian (orange) c) Monte carlo emission profile of a 6 x 11 µm hinge pixel d) Monte carlo emission profile of a 15 x 19 µm u-turn pixel e) Left, confocal image of a 3 mm biopsy punch from a rabbit transduced with ChRmine-mScarlet (red) and stained with RBPMS (green). Scalebar is 500 µm. Center, projected of RGC soma locations (blue) extracted from a rabbit retina and overlaid with the position of connected µLEDs in the hinge device (orange). Right, the same, but for a u-turn device. f) Number of RGCs that can be activated by at least one pixel of a hinge device (top) or u-turn device (bottom) versus axial distance from RGC layer (*x*-axis) or optical power (0.1 - 10 µW, see legend). Lines thickness indicates variation in the size of RGC activation field (soma and proximal dendrites, i.e. that area over which the mean irradiance determines action potential probability, 20-60 µm). g) Mean number of pixels that can activate an RGC that can be activated by at least one pixel for a hinge (top) or u-turn (bottom) device across a range of powers, axial distances, and activation fields as in (f). h) Fraction of pixels on a hinge (top) or u-turn (bottom) device capable of activating at least one RGC across a range of powers, axial distances, and activation fields as in (f). i) Number of addressable RGCs when all of the pixels of a hinge device (top) or u-turn (bottom) device are simultaneously activated across a range of powers, axial distances, and activation fields as in (f)

**Supplemental Figure 7:**
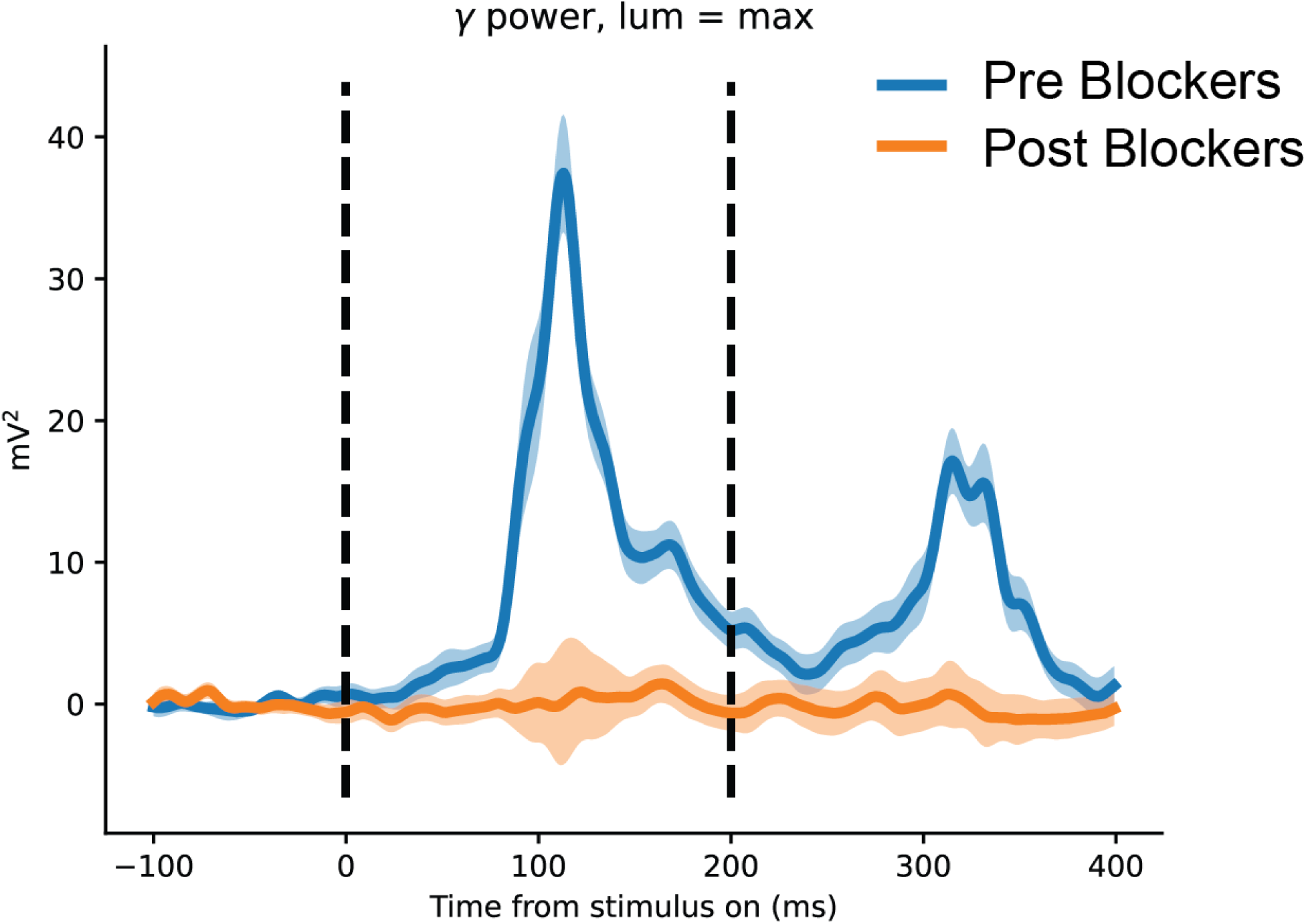
Intravitreal injection of synaptic blockers attenuates V1 VEPs in response to full-field visual stimulation with an LCD monitor (luminance). The blue trace shows average gamma power across the 32 channel grid implanted over V1 in response to the maximum luminance (641 lumens). The orange corresponds to gamma power in response to the same stimulation of the same eye 1 hour following injection of synaptic blockers (see methods). Shaded regions correspond to s.e.m. (n = 30 trials).

## Author Contributions

EK developed and performed rabbit visual cortex surgery, ECoG recording, and analyzed neural data

JB, TK, RS, and JMS performed eye surgery

KZ, JR, AR, KS, CW, AD and YF designed and built the FlexLED

SS and KR developed encapsulation for the device and developed the surgical carrier

MR and NS developed firmware, implanted electronics, and wireless power and data

AM performed whole cell patch clamp recordings

YLI, PD, ME, SC, KS, AR performed cell culture

AR, JB, PD, and KS performed histology

YLI and SM performed and analyzed sequencing experiments

JC and AM performed modeling experiments

AM, EK, KZ, and SM wrote the manuscript

AM, YK, MH conceived of and managed the project

## Acknowledgments

We thank E.J. Chichilinksy for advice and comments on the manuscript and Teresa Puthussery, Nicolas Pégard, Tim Gardner, and Morgan Brown for helpful advice and conversations.

